# Overcoming the inhibitory microenvironment surrounding oligodendrocyte progenitor cells following demyelination

**DOI:** 10.1101/2020.01.21.906073

**Authors:** Darpan Saraswat, Hani J. Shayya, Jessie J. Polanco, Ajai Tripathi, R. Ross Welliver, Suyog U. Pol, Richard A. Seidman, Jacqueline E. Broome, Melanie A. O’Bara, Toin H. van Kuppervelt, Joanna J. Phillips, Ranjan Dutta, Fraser J. Sim

## Abstract

Chronic demyelination in the human CNS is characterized by an inhibitory microenvironment that impairs recruitment and differentiation of oligodendrocyte progenitor cells (OPCs) leading to failed remyelination and axonal atrophy. By network-based transcriptomics, we identified sulfatase 2 (Sulf2) mRNA in activated human primary OPCs. Sulf2, an extracellular endosulfatase, modulates the signaling microenvironment by editing the pattern of sulfation on heparan sulfate proteoglycans. We found that Sulf2 was increased in demyelinating lesions in multiple sclerosis and was actively secreted by human OPCs. In experimental demyelination, elevated OPC Sulf1/2 expression directly impaired progenitor recruitment and subsequent generation of oligodendrocytes thereby limiting remyelination. Sulf1/2 potentiates the inhibitory microenvironment by promoting BMP and WNT signaling in OPCs. Importantly, pharmacological sulfatase inhibition using PI-88 accelerated oligodendrocyte recruitment and remyelination by blocking OPC-expressed sulfatases. Our findings define an important inhibitory role of Sulf1/2 and highlight the potential for modulation of the heparanome in the treatment of chronic demyelinating disease.

## INTRODUCTION

Endogenous myelin repair, known as remyelination, represents a regenerative process that can restore lost neurological function and prevent disease progression in degenerative conditions such as multiple sclerosis (MS) (Franklin, 2015). The failure of remyelination in MS has been attributed to a failure of oligodendrocyte progenitor cell (OPC) differentiation, in part, due to the presence of quiescent OPCs in regions of chronic demyelination (Wolswijk, 1998, Kuhlmann et al., 2008). Diverse signaling pathways have been identified that prevent timely OPC differentiation in rodent models and many small molecules have been described that improve the rate of murine remyelination by targeting individual rate-limiting steps (Franklin and Ffrench-Constant, 2017). Due to the complex nature of the lesion microenvironment, multiple pathways likely converge to prevent efficient remyelination. As such, we sought to define a therapeutic approach that would modulate the lesion microenvironment *en masse* to exact control over several inhibitory factors simultaneously to accelerate remyelination.

The accumulation of inhibitory extracellular matrix (ECM) components such as chondroitin sulfate proteoglycans and hyaluronan are known to impair remyelination in animal models and contribute to the inhibitory environment present in chronic demyelinated lesions in MS (reviewed in Harlow and Macklin, 2014, Sherman and Back, 2008, Pu et al., 2018). Recent analysis of MS GWAS studies identified numerous genes associated with modulation of CSPG deposition and extracellular matrix modification in general (Pu et al., 2020). Indeed, inhibition of CPSG synthesis by fluorosamine can prevent CPSG accumulation following demyelination and thereby accelerate myelin repair (Keough et al., 2016). In a similar manner, heparan sulfate proteoglycans (HSPGs) are deposited in both active and chronic demyelinating MS lesions (van Horssen et al., 2006). HSPGs represent a class of proteoglycans that interact with several growth factors (Viviano et al., 2004, Nawroth et al., 2007) and, thereby, have the potential to modulate several signaling cascades operant following demyelination. The regulation of signaling by HSPGs is highly dependent on modifications to the proteoglycan glycosaminoglycan sidechains. HSPG 6-O-sulfation (6S) occurs in a tissue and stage specific manner (Sarrazin et al., 2011) and can be edited by a family of two endosulfatases. Sulf1 and Sulf2 act specifically on the trisulfated IdoA2S-GlcNS6S disaccharide residue of HSPG, with no activity on chondroitin sulfate (Ai et al., 2006, Morimoto-Tomita et al., 2002). Sulfatase regulation of HSPGs is associated with alterations in bioavailability and activity of several growth factors and cytokines (reviewed in Rosen and Lemjabbar-Alaoui, 2010).

We previously identified a highly correlated module of genes whose expression was conserved across species and highly enriched among activated human OPCs (Pol et al., 2017). Among these genes, Sulf2 was among the mostly highly expressed and dynamically downregulated during differentiation. Herein, we describe sulfatase expression in adult OPCs following demyelination in mouse CNS and in demyelinated lesions in postmortem MS brain. In mouse, both Sulf1 and Sulf2 were expressed in tandem and largely restricted to oligodendrocyte lineage cells in adult CNS. Using genetic and pharmacological inhibition of sulfatases, we found that OPC-expressed sulfatases modulate their local microenvironment and facilitate HS sulfation-dependent WNT and BMP signaling. Furthermore, by conditional transgenic deletion of *Sulf1/2* in adult NG2-expressing OPCs, we demonstrate that sulfatases act to impair post-mitotic oligodendrocyte formation and inhibit remyelination. Taken together, our findings define a novel therapeutic target for the acceleration of OPC recruitment and differentiation, with potential translational applications in the treatment of demyelinating disease.

## MATERIALS AND METHODS

### Human CD140a/PDGFαR cell and tissue preparation

Fetal brain tissue samples, between 17 and 22 weeks of gestational age, were obtained from Advanced Bioscience Resources (Alameda, CA) with consent from patients under protocols approved by the University at Buffalo Research Subjects Institutional Review Board. Forebrain samples were minced and dissociated using papain and DNase as previously described (Conway et al., 2012). Magnetic sorting of CD140a/PDGFαR positive cells was performed as described (Pol et al., 2013). Human OPCs (hOPCs) were maintained on plates coated with poly-ornithine and laminin, in human neural differentiation (ND) media supplemented with 20 ng/ml PDGF-AA (PeproTech) and 5 ng/ml NT-3 (PeproTech) (Abiraman et al., 2015). Fetal tissue was fixed in 4% PFA and cryoprotected in a sucrose gradient (7.5% sucrose overnight, followed by 15% sucrose overnight), and frozen in OCT medium (Tissue-Tek). Serial 16 µm transverse sections were cut using a Leica cryostat and stored at -80°C.

### RNA extraction and RT-PCR analysis

Total RNA was isolated from hOPCs following mitogen withdrawal and real time RT-PCR performed as described in (Pol et al., 2017). Human SULF1, Fwd: ACCAGACAGCCTGTGAACAA and Rev: ATTCGAAGCTTGCCAGATGT, and SULF2, Fwd: TGAGGGAAGTCCGAGGTCAC and Rev: CTTGCGGAGTTTCTTCTTGC, specific primers were used to measure gene expression. Gene expression was calculated by normalizing to GAPDH and performing ΔΔC_t_ analysis.

### Cloning and lentiviral preparation

pLVTHM vectors containing shRNA targeting SULF2 (GGAGTGGTGGTGTCAATA) or a non-targeted scrambled control (AACAGTCGCGTTTGCGACTGG), were used as previously described (Nawroth et al., 2007). The pBARLHyg lentiviral reporter plasmid was a gift of Dr. Randall Moon (University of Washington) (Biechele and Moon, 2008). pGL3 BRE luciferase was a gift from Martine Roussel & Peter ten Dijke (Addgene plasmid # 45126) (Korchynskyi and ten Dijke, 2002). The lentiviral pTRIP-EF1a backbone was a gift from Abdel Benraiss (University of Rochester, Rochester, NY) (Sevin et al., 2006). To generate L-BRE-Luc, pGL3 BRE Luciferase was digested with KpnI (NEB), blunted using a commercial blunting kit (NEB) and digested with XbaI (NEB). The pTRIP-Ef1α plasmid was linearized by digestion with MluI (NEB), blunted, and digested with XbaI to remove Ef1α-mCherry-WPRE. The BRE-Luc fragment and the pTRIP-EF1a backbone were purified by gel extraction (Qiagen) and ligated with T4 DNA ligase (Invitrogen) to generate L-BRE-Luc. pBARLHyg, L-BRE-Luc and pLVTHM were packaged in lentivirus as previously described (Sim et al., 2006). Briefly, HEK 293T were triple transfected with viral vector and packaging plasmids pLP/VSVG (Invitrogen) and psPAX2 (AddGene), and viral supernatant was collected 42 hrs later. pLVTHM viral titer was determined by quantification of pLVTHM-driven GFP expression.

### Oligodendrocyte differentiation and immunocytochemistry

For knockdown studies, hOPCs were seeded at 5×10^4^ cells/mL and transduced with the indicated lentivirus after 24 hours. To initiate differentiation, PDGF-AA and NT-3 growth factors were removed, and cells cultured in ND media supplemented with human BMP7 (PeproTech), human WNT3a (R&D Systems) and/or PI-88 (gift of Medigen Biotechnology Corp, Taiwan), as designated, with media replenished after 48 hours. Four days following growth factor removal, cells were live stained with O4 IgM hybridoma supernatant (gift of Dr. James Goldman, Columbia University), fixed in 4% paraformaldehyde, and immunostained. For assessment of SULF2 expression, hOPCs were maintained as progenitors, fixed and stained for SULF2 (Abcam ab113405, 1:500). Where indicated, Brefeldin A (Cell Signaling Technology) was added to hOPCs at 5 µg/ml for five hours prior to fixation. Alexa 488, 594, and 647-conjugated secondary antibodies (Invitrogen) were used at 1:500 dilutions. Differentiation was quantified as the proportion of stained cells in 10 random fields at 10X magnification (using an Olympus IX70 microscope), representative of over 1,000 cells total in each condition.

### Assessment of hOPC HS sulfation by flow cytometry

hOPCs were cultured as previously described, trypsinized and collected in 0.02% EDTA/PBS. Cells were resuspended in HS3A8V (RB4CD12) phage display antibody (1:10, gift of Toin van Kuppevelt, Nijmegen Medical Center, Nijmegen, Netherlands).The His-tagged phage display antibody was detected by immunostaining with 6X-His Tag primary antibody (1:700, Abcam) and 1:500 Alexa-488 conjugated secondary antibody. All antibody incubations were for 30 minutes at +4°C. Flow cytometry was performed using a BD Fortessa flow cytometer. Dead cells were excluded by forward and side scatter-based gating, and doublet discrimination was performed. Fluorescence intensity (FITC-A) was quantified from ∼10,000 cells in each replicate, and data was normalized to peak fluorescence to facilitate presentation.

### Cell and secreted protein isolation and western blot

For cellular and secreted SULF2 expression analysis, hOPCs were cultured for 24 h in ND media supplemented with 20 ng/ml PDGF-AA and 5 ng/ml NT-3. For oligodendrocyte SULF2 expression, PDGF-AA and NT-3 was removed to allow oligodendrocyte differentiation for 3 days. Supernatants were collected, 0.22 μm filter sterilized and precipitated with acetone. Cells were washed in PBS, lysed and sonicated in a buffer containing 50 mM beta-glycerophosphate, 20mM HEPES buffer, 1% Triton-100, protease inhibitors (Roche), phosphatase inhibitor cocktail 2 (Sigma) and phosphatase inhibitor cocktail 3 (Sigma). For WNT and BMP pathway activity analysis, CG-4 rat OPCs were cultured on poly-D-lysine coated flasks in ND media supplemented with 10ng/mL PDGF-AA and 10ng/mL bFGF (PeproTech). Cells were treated with 50ng/mL human WNT3a, 50ng/mL human BMP7 and/or 100ug/mL PI-88 for the indicated time-points, washed in PBS, lysed and sonicated as above. Protein concentrations were determined using a Bradford assay (Bio-Rad), and 30µg protein was used for slot blot of secreted protein or added onto 10% poly-acrylamide gels for electrophoresis and western blot. Proteins were transferred to nitrocellulose membranes and blocked with commercial blocking buffer (Rockland) or 5% milk and 0.1% tween 20 (Calbiochem) in PBS depending on the primary antibody utilized. Primary antibodies against SULF2 (1:500, Abcam), phospho-SMAD 1/5 (1:1000, CST), active β-catenin (1:2000, Millipore), and β-actin (1:10,000, CST) were applied overnight at 4°C in blocking buffer. IR680RD and IR800CW secondary antibodies (Li-Cor) were diluted 1:5000 in Rockland blocking buffer. Blots were imaged using the Li-Cor Odyssey Infrared Imaging system.

### Luciferase Assays

OPCs were seeded at a density of 2.5 × 10^4^ cells/ml and maintained as progenitors in ND media supplemented with PDGF-AA and NT-3. One day post-seed, cells were infected with lentiviral luciferase reporter constructs for 24 hours, after which media was changed to fresh ND media supplemented with growth factors, BMP7, WNT and/or PI-88, as indicated. Twenty hours post-treatment, luminescence response was quantified using the Promega Bright-Glo reagent and a Bio-Tek plate reader, in accordance with manufacturer recommendations. Background luminescence was subtracted from all measurements and luminescence readings were normalized to the average value of the untreated control in each experiment. Data are presented as the mean ± SEM per individual human sample (three technical replicates).

### Animals and surgery

All experiments were performed according to protocols approved by the University at Buffalo’s Institutional Animal Care and Use Committee. NG2creER:Rosa26YFP animals were a gift from Akiko Nishiyama (University of Connecticut, Storrs, CT) (Zhu et al., 2011). *Sulf1/2* double-floxed (*Sulf1*^fl/fl^*Sulf2*^fl/fl^) animals were a gift from Dr. Xinping Yue (LSU Health Sciences Center, New Orleans, LA) (Ai et al., 2007). *Sulf1/2* double-floxed (*Sulf1*^fl/fl^*Sulf2*^fl/fl^) animals were crossed with the NG2creER; RosaYFP mice to generate three colonies NG2creER: YFP: Sulf1^fl/fI^, NG2creER: YFP: Sulf2^fl/fI^ and NG2creER: YFP: Sulf1/2^fl/fI^. Conditional *Sulf1, Sulf2 and Sulf1/2* knockout by cre-mediated recombination in adult NG2^+^ OPCs was achieved by intraperitoneal administration of tamoxifen (200 mg/kg, Sigma) for five days, the last of which occurred seven days prior to surgery. All experimental mice received tamoxifen in an identical manner and littermate controls lacking cre were compared with cre-containing cKO mice. For experiments using PI-88, eight-to eleven-week old female BalbC mice were purchased from Envigo.

Focal demyelination of the young adult (8-11 week) mouse spinal cord was induced as previously described (Welliver et al., 2018). Briefly, animals were anesthetized under isoflurane, and 0.5 μl, 1% lysolecithin (Lα-lysophosphatidylcholine, Sigma) was directly injected into the dorsal and ventral funiculus of the spinal cord between two thoracolumbar vertebrae. To assess the role of BMP and WNT modulation in the context of *Sulf1/2* deletion, CHIR-99021 (3µM, Axon MedChem), XAV939 (0.1µM, Tocris), A01 (100nM, Tocris) or LDN-193189 (100nM, Tocris) or matched volume saline (1:50) were co-injected with 1% lysolecithin (wt/vol). For experiments involving PI-88, 10 µg/ml PI-88 was co-injected with 1% lysolecithin, and demyelination induced by injection of the mixture into the spinal cord. Groups of WT and *Sulf1/2* cKO animals were co-injected with lysolecithin and drug/vehicle.

### Animal tissue processing and analysis

Animals were euthanized at 3, 5, 7, or 14 days post-lesion (dpl) by transcardial perfusion of saline followed by 4% paraformaldehyde under deep anesthesia. Tissue processing and lesion identification was performed as described previously (Welliver et al., 2018). Slides immediately adjacent to the lesion centers, identified by solochrome cyanine staining, were used for all immunohistochemical procedures. Primary antibodies utilized were rabbit anti-Olig2 (1:500, Millipore), mouse anti-CC1 (1:50, Millipore), mouse anti-GFAP (1:300, Sigma), rabbit anti-Iba1 (1:300, Wako Chemicals USA), cleaved caspase-3 (Asp175) (1:250, Cell signaling), and rabbit anti-NG2 (1:200, Millipore). Alexa 488, 594, and 647-conjugated secondary antibodies (Invitrogen) were used at 1:500. *In situ* hybridization for Plp1 was performed as described in (Welliver et al., 2018). Higher magnification examples of CC1/Olig2 staining demonstrating positive cell criteria used for quantification are shown in **Figure S9**. Images of spinal cords were captured at 20X magnification by wide field epifluorescence using an Olympus IX51 with Prior stage and were stitched together (FIJI). For each marker, the average from at least two sections was quantified for each animal. Lesions with cross-sectional area < 10,000 µm^2^ were excluded from the analysis, as were lesions which extended into surrounding grey matter. All quantification was performed by an investigator blinded to the identity of the samples.

### Electron microscopy

For assessment of remyelination, tissue was processed as described in Welliver et al. (2018). Briefly, mice were sacrificed at 14 day post lesion by transcardial perfusion with 2% glutaraldehyde in 0.1 M phosphate buffer and spinal cords were extracted (n = 5 per group), 1-mm-thick blocks surrounding the spinal cord lesion were processed through osmium tetroxide, dehydrated through ascending ethanol washes, and embedded in TAAB resin (TAAB Laboratories). One μm sections were cut, stained with 1% toluidine blue (Sigma-Aldrich), and examined by light microscopy to identify lesions. Selected blocks with lesions were trimmed, ultrathin sections cut, and examined by electron microscopy (Hitachi, HT7800). Images were acquired at 2500X magnification. Analyses of remyelinated axons and g-ratios were performed blinded. For remyelination counts, a minimum of 800 axons were counted for each animal from at least six different fields, with all animals per treatment group. Analysis of g-ratio was performed as described in Dillenburg et al. (2018). Briefly, axon and fiber diameter were measured using FIJI (diameter = 2 × √[area/π]). A minimum of 100 axons was analyzed per animal. The frequency distribution of axon diameter and g-ratio of remyelinated axons was calculated by binning on axon diameter and g-ratio.

### Human multiple sclerosis tissue

Multiple sclerosis tissue was prepared as described in Tripathi et al. (2019). Briefly, brains were collected as part of the tissue procurement program approved by the Cleveland Clinic Institutional Review Board. Brains were removed according to a rapid autopsy protocol, sliced (1 cm thick), and then fixed in 4% paraformaldehyde and 30 µm sections were cut for morphological studies. Lesions were characterized for demyelination by immunostaining using proteolipid protein (PLP). Chronic active demyelinated lesions were analyzed from two patients diagnosed with secondary progressive MS (53-57 yrs old, disease duration > 10 yrs, postmortem interval < 8 hrs).

### Fluorescent in situ hybridization (FISH)

Double label FISH was performed with probes targeting Sulf2 (GenBank: NM_001252578.1), Sulf1 (GenBank: NM_ 001198565.1), Plp1 (GenBank: NM_ 000533.4), Pdgfra1 (GenBank: NM_011058.2), Ki67 (GenBank: NM_001081117.2), Apcdd1 (GenBank: NM_133037.3), ID4 (GenBank: NM_031166.2) mRNA by using the RNAscope fluorescent multiplex detection kit (Advanced Cell Diagnostics) according to the manufacturer’s instructions. Postnatal day 7 and 28, and 24-week old mice were euthanized and perfused with 0.9% saline followed by 4% paraformaldehyde. Dissected mice brains and spinal cords and cryopreserved in sucrose gradient (10, 20, 30%) for 24hr and were snap frozen in OCT and sectioned coronally at 16 µm thickness. Fixed sections were baked at 60°C for 1 hr, washed with ethanol, followed by tissue pretreatment, probe hybridization following RNAscope fluorescence multiplex assay and sections were counterstained with DAPI to visualize nuclei. Positive signals were identified as punctate dots present in nucleus and cytoplasm. All tissues were tested using negative control probes (ACD Bio) to control for non-specific binding (**Figure S2a**). For Sulf1 and Sulf2 probes, the expression patterns were verified by comparison with known sulfatase-specific patterns of neuronal expression in cortical layer V (Sulf2)(Zeisel et al., 2015) and layer VI (Sulf1)(Allen Brain Atlas) (**Figure S2b-f**). For all RNAscope-based imaging, confocal Z stack images with optical thickness 0.2 µm were taken and stacked images are shown (Zeiss LSM 510 Meta confocal).

### Statistical analyses

All quantification and data analysis were performed by an investigator blinded to the identity of the samples. All statistical analyses were performed using GraphPad Prism (San Diego, CA). Data were compared by Student’s *t* test, one-way ANOVA, or two-way ANOVA, where appropriate; significance was considered at p<0.05. Each figure legend contains the statistical test and significance values. Data are reported as mean ± SEM. The data that support the findings of this study are available from the corresponding author upon reasonable request.

## RESULTS

### Heparan sulfate proteoglycan 6-O endosulfatases are highly expressed by OPCs

Transcriptomic network analysis of human OPC (hOPC) differentiation identified a module of highly correlated and species conserved genes associated with progenitor fate (Pol et al., 2017). Among these high connected genes, heparan sulfate endosulfatase 2 (SULF2) was identified in the OPC-associated module (M5). Real time RT-PCR analysis confirmed profound down-regulation during hOPC *in vitro* differentiation (**Fig. 1a**). Expression of SULF2 was relatively restricted to OPCs as SULF2 mRNA expression was >14-fold higher in primary human PDGFαR^+^ OPCs than CD133^+^ neural progenitor cells (Wang et al., 2013). Likewise, SULF2 expression was maintained in oligodendrocyte-biased PDGFαR^+^O4^+^ progenitors (Abiraman et al., 2015) and adult human A2B5^+^ OPCs but not enriched in adult human astrocyte or microglial isolates (Sim et al., 2009).

**Figure 1.**
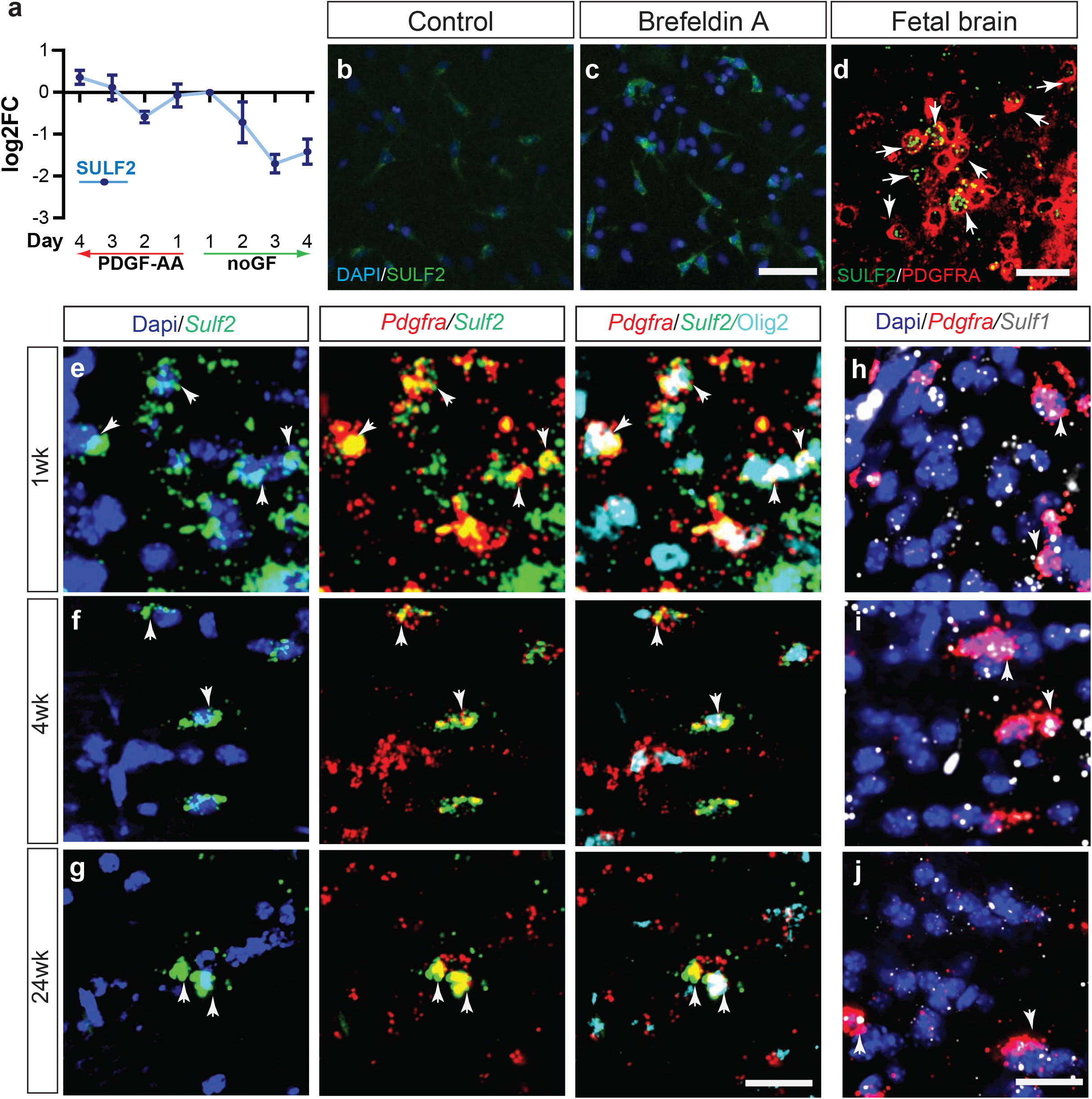
Sulfatase is expressed by mouse and human OPCs. **a**, RT-PCR analysis of SULF2 mRNA in human primary OPCs during *in vitro* differentiation after PDGF-AA removal. **b**-**c**, SULF2 protein is expressed at low levels in hOPCs (**b**), and immunoreactivity substantially increased after treatment with the protein secretion inhibitor brefeldin A (5 µg/ml) (**c**). Confocal micrograph of SULF2 mRNA expressing PDGFRA^+^ OPCs in fetal human brain (**d**). **e-g**, *Sulf1* and *Sulf2* expression was analyzed by confocal microscopy in mouse corpus callosum during normal development by RNAscope *in situ* hybridization and combined with Olig2 Immunohistochemistry (IHC) at postnatal day 7 (**e, h**), day 28 (**f, i**) and at 24 weeks (**g, j**). *Sulf1* mRNA (cyan), *Sulf2* mRNA (green), *Pdgfra* mRNA (red) and Olig2 protein (pink). White arrows denote *Pdgfra*^*+*^ OPCs expressing *Sulf1* and *Sulf2*. Arrowheads denote *Sulf1*^*+*^*Pdgfra*^*+*^ OPCs. Yellow arrows denote Olig2^+^*Pdgfra*^*-*^ oligodendrocytes that express *Sulf2*. Scale: 100 µm (**b-c**), 20 µm (**d)** and **20** µm (**e-j)**.

As heparan sulfate function has not been extensively studied during OPC development, we examined the expression of genes involved in HS biosynthesis, post-synthetic modifications of HS, and HSPG core proteins in mouse development (**Fig. S1**). We observed that several HS-associated genes were down-regulated with differentiation. Down-regulation of HS biosynthetic genes during OPC differentiation was consistent with previous studies that suggested reduced heparan/HS abundance during OPC differentiation (Stringer et al., 1999, Properzi et al., 2008). Together, these findings suggest that the heparanome is dynamically regulated in the progenitor state and becomes more static as OPCs differentiate to mature OLs. Remarkably, both mouse *Sulf1* and *Sulf2* mRNAs were almost entirely restricted to the oligodendrocyte lineage (**Table S1**). In contrast to mouse, RNA-seq of primary hOPCs revealed very high levels of *SULF2* expression (∼100 FPKM) whereas *SULF1* mRNA almost undetectable (<0.5 FPKM) (**Table S1**). SULF2 protein expression was readily detected in hOPCs (**Fig. 1b**). Consistent with active secretion of sulfatases (Morimoto-Tomita et al., 2002, Tang and Rosen, 2009), blockade of the secretory pathway with Brefeldin A led to cytoplasmic SULF2 protein accumulation in hOPCs (**Fig. 1c**).

We observed high expression of *SULF2* mRNA *in vivo* restricted to a subset of *PDGFRA*^*+*^ OPCs located in the human subventricular zone and white matter of 22-week fetal brain (**Fig. 1d**). In 1-week old mouse brain, almost all *Pdgfra*^*+*^ OPCs expressed *Sulf2* mRNA (**Fig. 1e and Fig. S2b**) while *Sulf1* mRNA was restricted to a subset of OPCs (**Fig. 1h and Fig. S2b**).During early postnatal development both sulfatases were also detected in immature oligodendrocytes but expression in oligodendrocytes was not sustained in the adult (**Fig. 1f-j**). Consistent with previous reports (Zeisel et al., 2015)(Allen Brain Atlas), Sulf1 and Sulf2 mRNAs were highly expressed within specific cortical layers **(Fig. S2c-f)**. Sulf2 mRNA was also expressed by a small subset of Gfap^+^ astrocytes and Iba1^+^ microglial cells **(Fig. S2g-j)**. In the corpus callosum of aged adult mice (24-week), only *Pdgfra*^*+*^ OPCs retained high levels of *Sulf1/Sulf2* mRNA (**Fig. 1g/j**). Combined with our expression data in human cell isolates, these findings suggest that OPCs expressed *Sulf1*/*Sulf2* throughout developmental and into adulthood consistent with a functional role in OPC homeostasis.

### Sulfatase inhibits oligodendrocyte production following demyelination

We examined the pattern of sulfatase expression following demyelination by inducing demyelination in adult mouse spinal cord by focal injection of lysolecithin. We observed increased expression of *Sulf1* and *Sulf2* mRNA within the lesion at 5 days post-lesion (5 dpl) (**Fig. 2a** and **b**). At this time-point, OPCs are actively recruited into the lesion. We observed that many *Pdgfra*^*+*^ OPCs within the lesion expressed *Sulf1* or *Sulf2* mRNAs (**Fig. 2a-i**), while a subset expressed both *Sulf1* and *Sulf2* (**Fig. 2e**,**j**). As in normal brain, *Sulf2* expression was restricted to OPCs in the unlesioned spinal cord white matter. In contrast, following demyelination *Sulf2* was also expressed by a very small subset of Gfap^+^ astrocytes and Iba1^+^ microglia in the vicinity of and within the lesion while only very few *Sulf1*^+^ astrocytes were observed (**Fig. S3**). Of note, Pdgfrb^+^ pericytes did not express detectable Sulf1 or Sulf2 following demyelination. These observations suggest that both sulfatases are up regulated by OPCs following demyelination and that sulfatase expression was largely restricted to OPCs and oligodendrocyte lineage cells.

**Figure 2.**
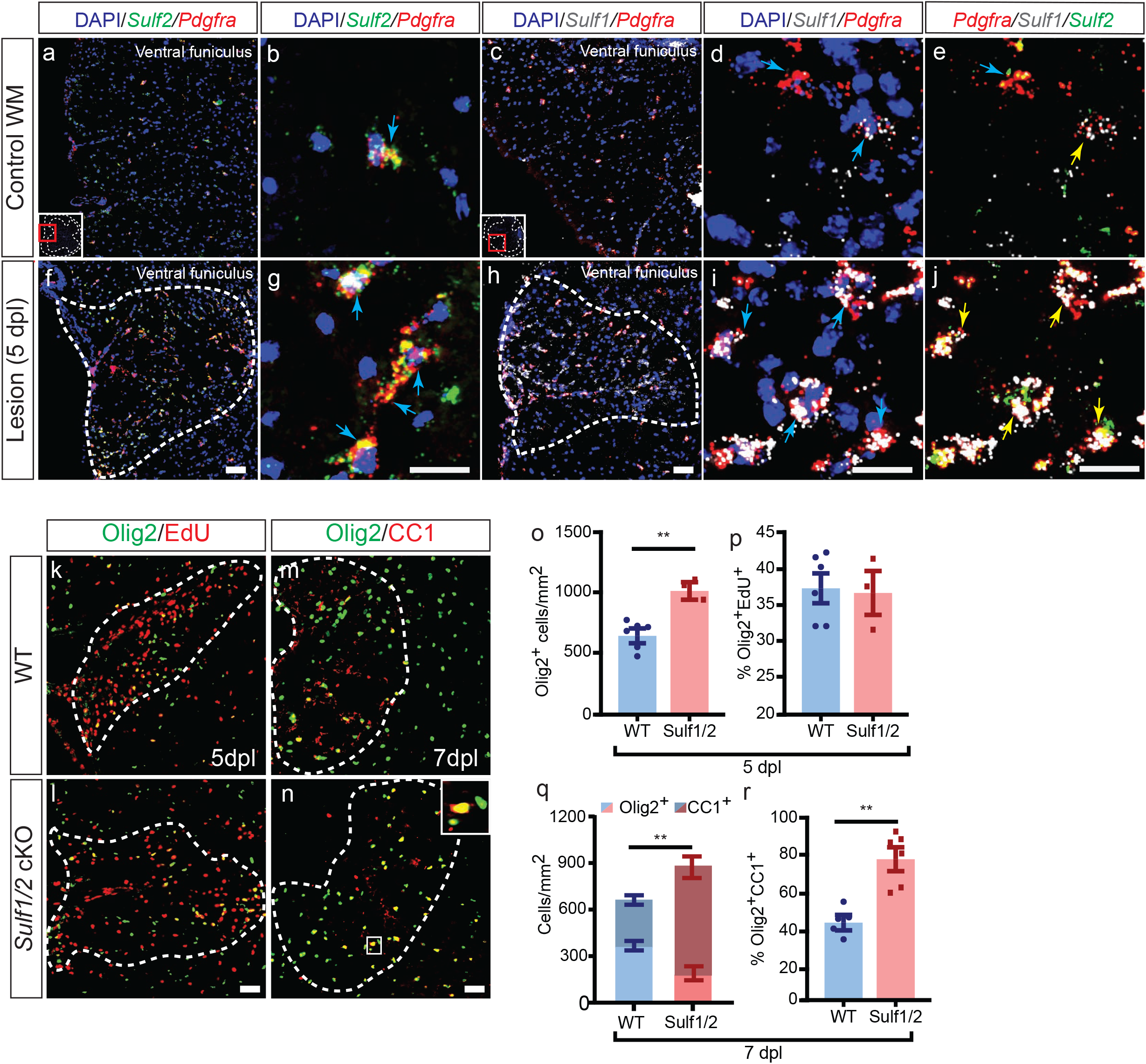
Conditional genetic ablation of OPC-expressed *Sulf1/2* accelerates oligodendrocyte recruitment and lesion colonization following demyelination. **a-j**, Dual RNAscope *in situ* hybridization (ISH) of mouse spinal cord (**a-e**) revealed increased *Sulf1* and *Sulf2* mRNA expression following demyelination (**f-j**; 5 dpl). *Sulf1* (cyan), *Sulf2* (green), or *Pdgfra* mRNA (red) and Olig2 protein (pink). Yellow arrows denote *Pdgfra*^*+*^ OPCs that co-expresses *Sulf1/2*. Blue arrows denote *Sulf2*^*+*^*Pdgfra*^*+*^ OPCs or *Sulf1*^*+*^*Pdgfra*^*+*^ OPCs. RNAscope images were imaged by confocal microscopy, others using wide field epifluorescence. Inducible NG2creER-mediated OPC-specific ablation of *Sulf1*/*2* (Sulf1/2 cKO) in adult mice followed by lysolecithin-induced demyelination (**k-r**). Analysis of OPC dynamics at 5 days post-lesion (dpl) (**o-p**) and 7 dpl (**q-r**). At 5 dpl, Olig2^+^ cell density was significantly increased in *Sulf1/2* cKO (**o**). No difference in the proportion of proliferating EdU^+^Olig2^+^ cells compared to wild type (**p**) (** p < 0.01, t-test, n= 3 - 6 per group). At 7 dpl, *Sulf1/2* cKO exhibited increased overall Olig2^+^ and post-mitotic Olig2^+^CC1^+^ oligodendrocyte density (**m-n**) (cells/mm^2^ quantified in **q**, ** p < 0.01). The percentage of CC1^+^Olig2^+^ oligodendrocytes among Olig2^+^ cells was increased in *Sulf1/2* cKO animals compared to controls (**r**, *** p < 0.001, n = 3 - 6 per group). Mean ± SEM shown. Scale: 20 µm (**a**,**f**,**c**,**h**), 40 µm (**b**,**g**,**d-j**), and 10 µm (**k-n)**.

To determine the functional role of OPC-expressed sulfatases following demyelination, we performed tamoxifen-induced conditional ablation of *Sulf1/Sulf2* 1-week prior to lysolecithin-induced demyelination. In order to ablate *Sulf1/2* specifically from adult OPCs, we crossed *NG2creER* mice (Zhu et al., 2011) with floxed *Sulf1/Sulf2* mice (Ai et al., 2007). Littermate controls lacking cre-expression were also injected with tamoxifen. At 5 dpl, we observed a significant increase in Olig2^+^ oligodendrocyte lineage cell density in *Sulf1/2* cKO mice compared to wildtype (wt) (n = 3-6 animals per group, t-test p < 0.01) (**Fig. 2k-l, o**). To assess OPC proliferation animals were terminally injected with EdU. Increased Olig2^+^ cell density was not due to increased proliferation (EdU%, p = 0.85) (**Fig. 2p**). This suggested that ablation of sulfatase encourages a favorable microenvironment for OPC migration and recruitment.

Significantly increased density of Olig2^+^ oligodendrocyte lineage cells was maintained at 7 dpl (**Fig. 2m-n**). At 7 dpl, *Sulf1/2* cKO mice exhibited a 37% increase in density of Olig2^+^ cells (n = 6-8 animals per group, t-test p < 0.01) (**Fig. 2q**). Accompanying the increase in total oligodendrocyte lineages cells, we noted a 73% increase in the proportion of CC1^+^Olig2^+^ amongst Olig2^+^ cells in *Sulf1/2* cKO relative to wt controls (p < 0.001) (**Fig. 2r**). This resulted in a 2.2-fold increase in the density of Olig2^+^CC1^+^ post-mitotic oligodendrocytes (p < 0.001). Intriguingly, the increase in oligodendrocyte differentiation as this stage resulted in the relative depletion of Olig2^+^CC1^-^-defined cells at this stage.

To more directly define progenitor recruitment following demyelination, we assessed the density of *Pdgfra* mRNA-defined OPCs and immature oligodendrocytes at each stage following demyelination and determined the rate of progenitor proliferation by co-labeling *Pdgfra*^*+*^ OPCs with Ki67 (**Fig. S5**). As shown previously (Sim et al., 2002), in wildtype animals Pdgfra^+^ OPC density was greatest at 5 dpl and gradually declined thereafter (**Fig. S5d**). Following *Sulf1/2* cKO, we observed significantly increased *Pdgfra*^*+*^ OPC density at both 5 and 7dpl compared to matched wt controls (Holm-Sidak multiple comparison test, n = 4-6 animals, p < 0.01) (**Fig. S5d**). By 14dpl, the *Sulf1/*2-dependent increase in OPC density was no longer present (n= 3-4 per group, p =0.64). The proportion of dividing Ki67^+^ OPCs likewise decreased over the same time period. Consistent with the EdU analysis above (**Fig. 2P**), *Sulf1/2* cKO did not influence the proportion of Ki67^+^Pdgfra^+^ OPCs at any stage following demyelination suggesting that proliferation of OPCs was not effected by *Sulf1/2* ablation (*Ki67*%, p = 0.99) (**Fig. S5e**). Next to determine, whether differential cell death contributed to the differences observed in oligodendrocyte lineage recruitment, we analyzed apoptosis via cleaved caspase 3 within the Olig2^+^ cell population (**Fig. S5f-h)**. The differences in proportion of apoptotic OPCs between *Sulf1/2* cKO and wt mice did not reach significance (n = 3-4, p = 0.28)(**Fig. S5h**) and, as such, likely do not account for the large differences in Olig2^+^ cell density observed (**Fig. 2o and q)**. Thus, sulfatases *Sulf1/2* act to impair recruitment and early differentiation of OPCs following demyelination.

### Individual sulfatase deletion is sufficient to improve oligodendrocyte generation following demyelination

In order to define whether the effects of *Sulf1/2* cKO was mediated specifically via deletion of either *Sulf1* or *Sulf2*, we assessed the effect of individual *Sulf1/2* cKO on OPC recruitment and differentiation following demyelination at 7 dpl (n = 5-6 animals per genotype) (**Fig.3a-g**). Interestingly, individual *Sulf2* deletion resulted in a significant increase in the density of Olig2^+^ oligodendrocyte lineage cells that resembled that of *Sulf1/2* double cKOs (**Fig. 3e**). In contrast, *Sulf1* deletion resulted in only a relatively mild increase to Olig2^+^ density relative to control (**Fig. 3e**). Indeed, two-way ANOVA treating *Sulf1/2* as dependent variables indicated a main effect of *Sulf2* but not *Sulf1* genotype on Olig2^+^ recruitment (*Sulf2* p < 0.0001, *Sulf2* p = 0.67). Consistent with a shared mechanism, a significant interaction between *Sulf1* and *Sulf2* was found (F (1, 17) = 17.4, p = 0.001) (**Fig. 3e**). In contrast, individual deletion of either *Sulf1* or *Sulf2* resulted in a highly significant increase in CC1^+^Olig2^+^ oligodendrocyte density relative to WT (two-way ANOVA, main effects p < 0.05). A highly significant interaction of *Sulf1* and *Sulf2* genotype effect on Olig2^+^CC1^+^ cell density was also consistent with a common function of either gene (F (1, 17) = 17.4, interaction p = 0.0006) **(Fig. 3f)**. Likewise, as an estimate of differentiation, the proportion of CC1^+^Olig2^+^ cells among Olig2^+^ oligodendrocyte lineage cells was increased following ablation of either *Sulf1* or *Sulf2*, and comparable with *Sulf1/2* double cKO mice (F (1, 16) > 10, p < 0.01 main effects and interaction) (**Fig. 3g**).

**Figure 3.**
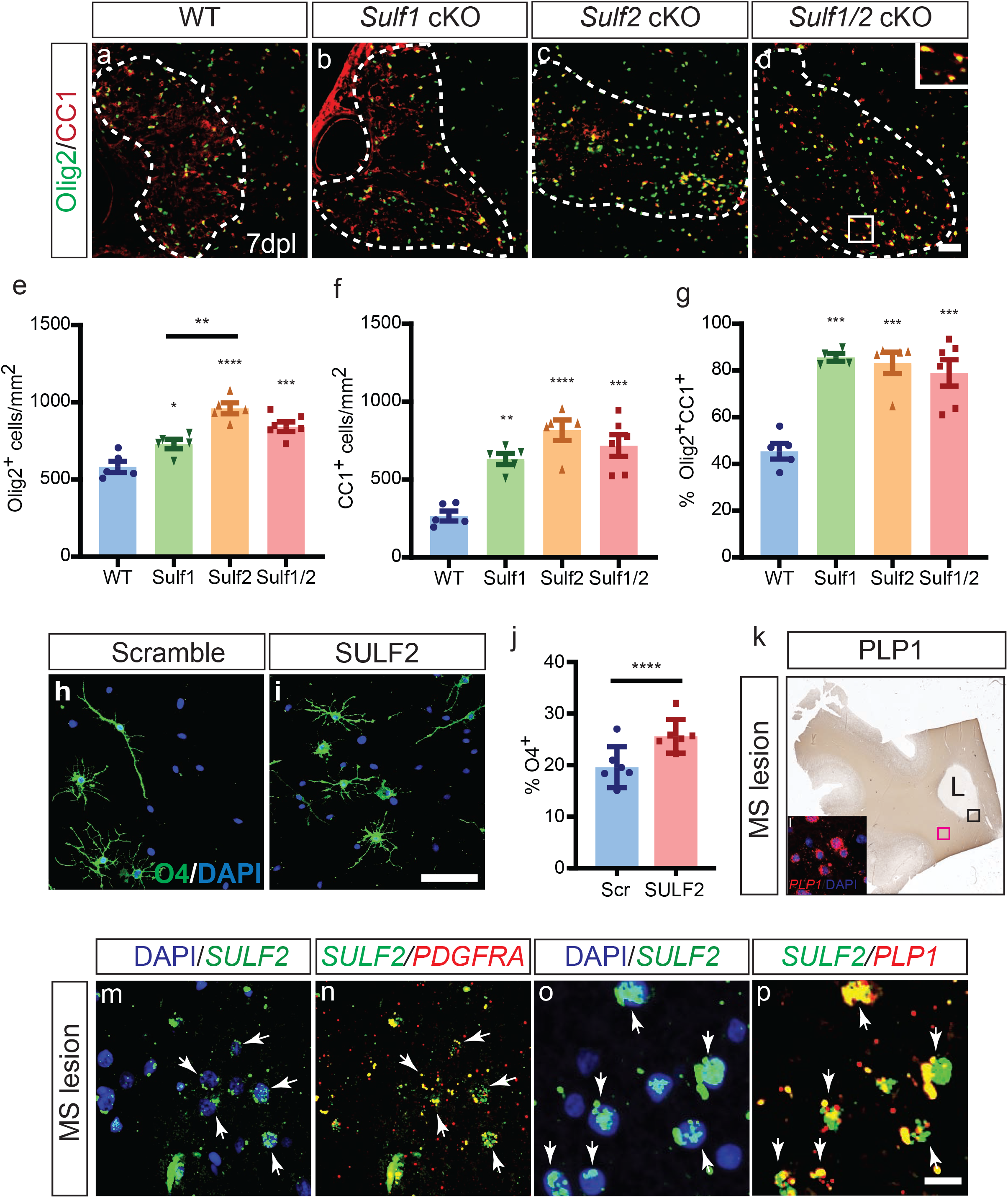
Sulf2 deletion results in improved oligodendrocyte recruitment and differentiation and is up regulated in chronic active MS lesions. Tamoxifen-dependent OPC-specific ablation of either *Sulf1, Sulf2* alone or both *Sulf1/2* in NG2^+^ OPCs was initiated prior to demyelination. Control animals (lacking cre) were treated in an identical manner. Widefield immunofluorescence for Olig2 and CC1 at 7 dpl (**a-d**). The density (cells/mm^2^) of Olig2^+^ oligodendrocytes lineage cells (**e**), Olig2^+^CC1^+^ post-mitotic oligodendrocytes (**f**) and percentage of CC1^+^ oligodendrocytes (**g**) was quantified (n = 5 - 6 mice per group). Two-way ANOVA for Sulf1 and Sulf2 genotypes. Holm-Sidak post-test vs. wild-type control *, **, ***, *** indicates p < 0.05, 0.01, 0.001, 0.0001 respectively. **h-j**, human primary OPCs infected with lentiviral SULF2 knockdown (KD) or scrambled control cultured in mitogen-free conditions for 4 days. Oligodendrocyte differentiation assessed by O4 immunocytochemistry (green) and DAPI (blue) (**h-i**). **j**, SULF2 KD accelerated O4^+^ oligodendrocyte differentiation (**** p < 0.0001 paired t-test, n=6 fetal human samples). **k**, PLP1 immunohistochemistry of chronic active human MS lesion and surrounding normal white matter (NAWM). **l** (insert in K), RNAscope for *PLP1* mRNA in NAWM (pink box in k). **m-p**, Confocal-based RNAscope *in situ* hybridization at lesion border (black box in k) for *SULF2* (green), *PDGFRA* (red, **n**) and *PLP1* (red, **p**) mRNA. Arrow represents colocalization of *SULF2* with *PDGFRA* and *PLP1* mRNAs. Mean ± SEM is shown. Scale: 20 µm (**a-d, n-q**), and 50 µm (**i-j**).

Together, these data suggest that *Sulf1* and *Sulf2* both contribute to delayed OPC recruitment and oligodendrocyte differentiation following demyelination. As deletion of either sulfatase resulted in substantial alterations in the density of oligodendrocytes and oligodendrocyte lineage cells, it is clear that the loss of either *Sulf1* or *Sulf2* was not fully compensated by the remaining sulfatase gene.

### *SULF2* expression inhibits hOPC differentiation and is enriched in MS lesions

Having established that *Sulf2* can impair OPC recruitment and differentiation in mice following demyelination, we next studied the functional role of sulfatase expression in purified human OPCs. RNA-seq and qPCR revealed that SULF2 was the principal sulfatase expressed by hOPCs (**Table S1 and Fig. S4a**). We examined the function of SULF2 in hOPCs following infection with lentivirus expressing SULF2 shRNA (Phillips et al., 2012), which significantly reduced *SULF2* mRNA expression by 72.9 ± 5.7% (**Fig. S4b**) (p < 0.0001, n = 8 individual fetal samples). Importantly, *SULF1* mRNA expression was not increased in compensation following SULF2 knockdown (**Fig. S4c**) (p = 0.90) and remained at the lower detection limit (C_t_ > 30). Importantly, SULF2 knockdown significantly accelerated human O4^+^ oligodendroglial differentiation by 46% compared to scrambled controls (**Fig. 3i-j**) (n = 6 samples, p < 0.0001). To determine if hOPCs actively secreted SULF2, we performed slot blot analysis of conditioned media as well as cell lysates (**Fig. S4d**). Consistent with *SULF2* mRNA downregulation upon differentiation (**Fig. 1a**), both cellular and secreted SULF2 protein expression decreased during hOPC differentiation.

As SULF2 acted to prevent efficient hOPC differentiation *in vitro*, we next asked whether *SULF2* was expressed by OPCs in adult human brain and in chronic active demyelinated lesions from secondary progressive MS patients (n = 2). SULF2 mRNA was expressed by *PDGFRA*^*+*^ OPCs in normal appearing white matter and substantially increased around the demyelinated lesion border (**Fig. 3l-p**). We confirmed that these *SULF2* mRNA transcripts were restricted to OPC and oligodendrocyte cells by their colocalization with *PDGFRA* (**Fig. 3m-n**) and *PLP1* transcripts, respectively (**Fig 3o-p**). Almost all *PDGFRA*^*+*^ and *PLP1*^*+*^ cells observed at the lesion border expressed SULF2 mRNA. Together, these data suggest that SULF2 is up regulated in a similar manner following demyelination in the human brain and *in vitro* act in a similar manner to prevent efficient OPC differentiation.

### Conditional sulfatase ablation accelerates remyelination following demyelination

To address whether Sulf1/2 modulation of HSPG sulfation could influence oligodendrocyte density and remyelination following lysolecithin-induced demyelination, additional cohorts of *Sulf1/2* cKO and control mice were sacrificed at 14 dpl. As observed at 5 and 7 dpl, demyelinated lesions were similar in size in *Sulf1/2* cKO and control mice (n = 4, t-test p = 0.81) (**Fig. 4a, b**). Sulfatase deletion resulted in a 2-fold increase in the density of *Plp1*^+^ oligodendrocytes and 2.6-fold increase in Olig2^+^CC1^+^ oligodendrocyte density (n = 4 per group, p < 0.05) (**Fig. 4c-d, k, l**) as seen at 7 dpl. Likewise, increased mature oligodendrocyte density in *Sulf1/2* cKO animals was accompanied by increased total density of Olig2^+^ oligodendrocyte lineage cells relative to wt controls (p< 0.01) (**Fig. 4l**). The density of Olig2^+^CC1^-^ defined cells was equivalent between groups at 14 dpl (**Fig. 4l**) as was the density of Pdgfra^+^ OPCs (**Fig. S5d**). The percentage of Olig2^+^CC1^+^ oligodendrocytes within the Olig2^+^ population was no longer enhanced at 14 dpl (p = 0.72) (**Fig. 4m**). Taken together, *Sulf1/2* cKO resulted in accelerated OPC recruitment and differentiation of new oligodendrocytes leading to increased density of both mature oligodendrocytes and OPCs following demyelination in the CNS.

**Figure 4.**
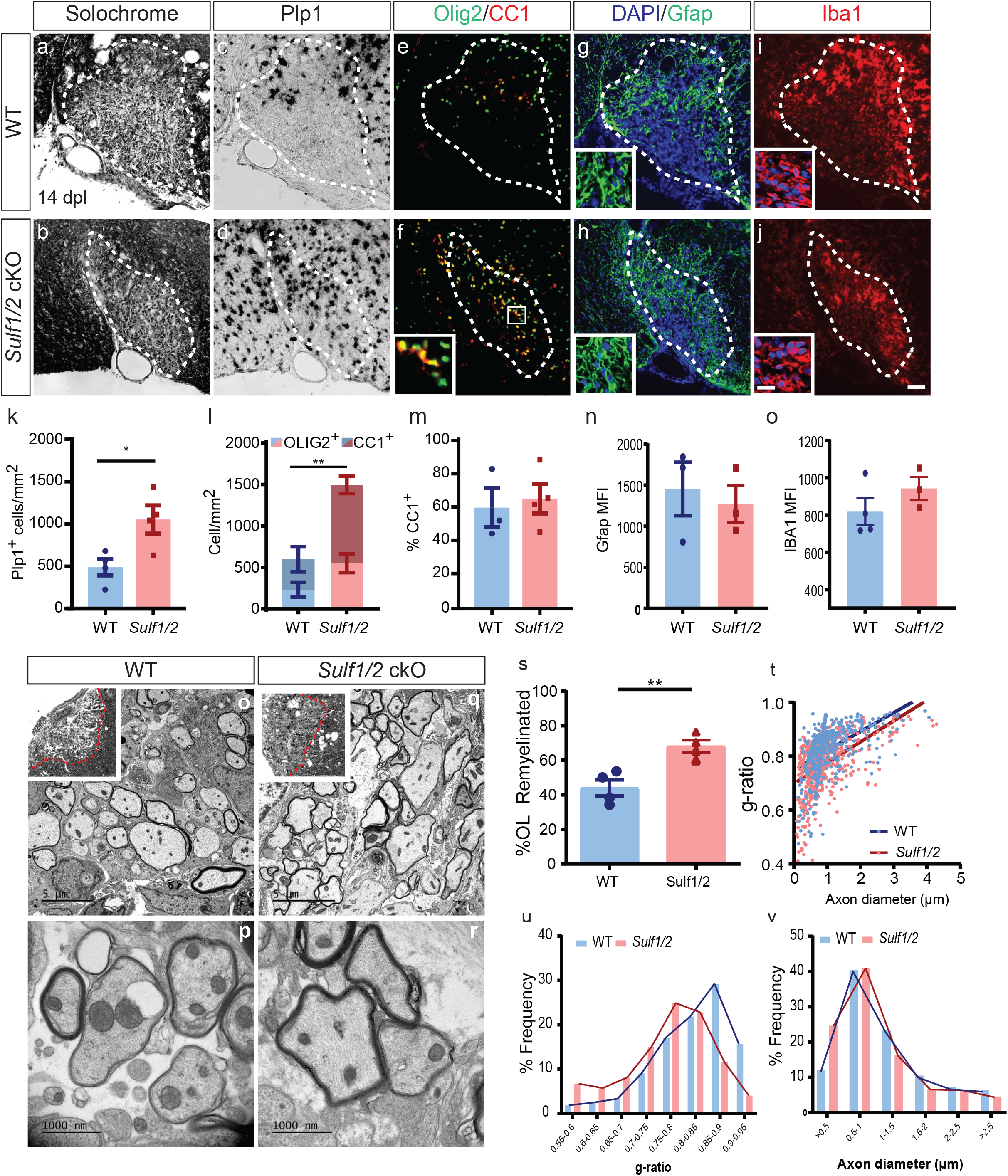
Conditional ablation of *Sulf1/2* accelerates remyelination. **a-b**, Solochrome cyanine stained lesions at 14 dpl. Oligodendrocyte differentiation was assessed by *Plp1* ISH (**c-d**) and Olig2^+^CC1^+^ immunofluorescence (**e-f**). Astrogliosis was assessed by GFAP and image insert shows higher magnification image to show morphology (**g-h**), and the microglial responses assessed by Iba1 and image insert shows higher magnification image to show morphology (**i-j**). Quantification of *Plp1*^+^ oligodendrocyte cell density (**k**), Olig2^+^ and CC1^+^ density (**l**) and percentage of CC1^+^ cells within the Olig2 population (**m**) (n = 3 - 4 mice per group). * indicates t-test p<0.05.**n**, Mean fluorescence intensity (MFI) of GFAP and IBA1 cells in lesion. **o-v**, Analysis of remyelination by EM at 14 dpl. **o**,**q**, Semithins inserts showing lesions in ventrolateral white matter. **s**, Proportion of remyelinated axons (** indicates t-test p < 0.01), **t**, relationship between axon diameter and *g*-ratio (linear regression shown). Frequency distribution of axonal *g*-ratio (**u**) and axon diameter (**v**) in lesion (n = 4 animals/group, ≥400 axons). Mean ± SEM is shown. Scale: 20 μm (**a-j**) and 5 μm (**o, q**) and 1000 nm (**p, r**).

As OPC secreted sulfatases may regulate local signaling in other cell types, we examined the effect of OPC-specific deletion of *Sulf1/2* on astrocyte and microglial activation following demyelination. We did not observe differences in the gross pattern or intensity of astrocytic Gfap immunoreactivity (p = 0.66) (**Fig. 4g, h**). Likewise, the overall distribution and intensity Iba1^+^ microglia/macrophages staining was not significantly altered by *Sulf1/2* cKO (p = 0.26) (**Fig.4i-j, n**). Thus, while paracrine effects of OPC-expressed *Sulf1/2* cannot be excluded, there were no overt effects on the gliotic response following demyelination.

We next investigated whether decreased sulfatase activity in OPCs might accelerate remyelination (**Fig. 4o-v**). At 14 dpl, *Sulf1/2* cKO significantly increased the proportion of oligodendrocyte remyelinated axons relative to WT controls (n = 4 animals per group, t-test p = 0.01) **(Fig. 4s)**. While a small proportion of axons were remyelinated by Schwann cells, we did not observe any changes in the proportion of Schwann cell myelinated axons following *Sulf1/2* cKO (data not shown). Consistent with an acceleration of oligodendrocyte differentiation, myelin thickness was significantly increased in remyelinated axons following *Sulf1/2* deletion. The myelin *g*-ratio, which represents the ratio of axon to axon plus myelin diameter, was significantly decreased in *Sulf1/2* cKO (*g*-ratio calculated from n = 4 animals, t-test p < 0.05). Linear regression analysis of individual *g*-ratio vs. axon diameter also demonstrated a significant effect of *Sulf1/2* cKO (**Fig. 4t**). Likewise, the distribution of *g*-ratios demonstrated a significant reduction in the frequency of very thinly remyelinated axons consistent with improved remyelination initiation following Sulf1/2 cKO (Two-way ANOVA, Sidak post hoc, 0.85-0.9 p < 0.0001 and 0.9-0.95 p<0.01) (**Fig. 4u**). *Sulf1/2* cKO did not influence the distribution of axonal diameter within the lesion suggesting that deletion of *Sulf1/2* does not induce axonal swelling (**Fig. 4v**). These results indicate that OPC-expressed sulfatases act as negative regulators for remyelination in a cell-autonomous manner.

### WNT/BMP signaling inhibits oligodendrocyte differentiation following demyelination in a sulfatase-dependent manner

Pathological activation of both BMP and WNT signaling prevents efficient remyelination by inhibiting OPC differentiation (Deininger et al., 1995, Ara et al., 2008, Fancy et al., 2009). Importantly, antagonism of either WNT or BMP signaling can accelerate OPC differentiation and remyelination (Fancy et al., 2011, Sabo et al., 2011). As sulfatase ablation has been shown to impair both BMP (Otsuki et al., 2010) and WNT signaling (Nawroth et al., 2007), we hypothesized that *Sulf1/2* cKO increased oligodendrocyte density due to modulation of WNT and BMP signaling in OPCs. To this end, we used pharmacological approaches to specifically activate or inactivate either WNT or BMP signaling in the context of *Sulf1/2* deletion at 7 dpl **(Fig. 5a-m)**. Consistent with previous studies and data presented above, two-way ANOVA revealed significant effects of drug treatment and *Sulf1/2* genotype on all parameters as well as highly significant interaction between them (n = 3-8 animals per group) (**Tables S2 and S3**).

**Figure 5.**
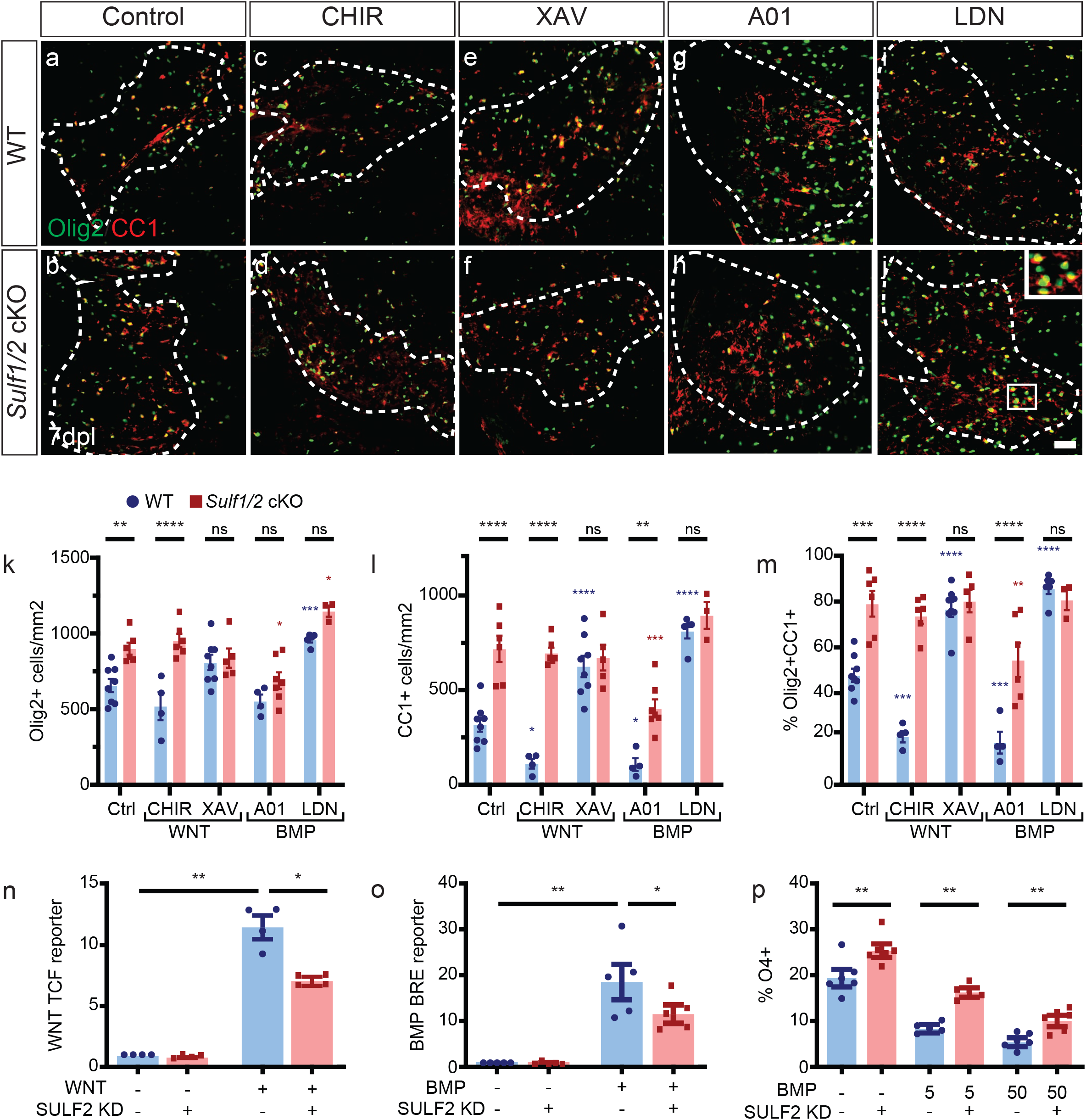
Sulfatase promotes inhibitory WNT and BMP signaling on oligodendrocyte differentiation following demyelination. Pharmacological manipulation of WNT and BMP signaling was performed in the context of Sulf*1/2* cKO. WNT pathway activator (CHIR-99021, 3 µM), WNT antagonist (XAV939, 100nM), BMP pathway activator (A01, 100nM) or receptor antagonist (LDN-193189, 100nM) were co-injected with lysolecithin. **a-j**, Oligodendrocyte lineage cell density was assessed by Olig2 (green) and CC1 (red) widefield immunofluorescence. Density of Olig2^+^ oligodendrocyte lineage cells (**k**), Olig2^+^CC1^+^ post-mitotic oligodendrocyte (**l**), and percentage of CC1+ oligodendrocytes (**m**) quantified (n = 3 - 8 group, mean ± SEM). Two-way ANOVA performed. Holm-Sidak multiple comparisons tests vs. ctrl wild-type (blue *) or ctrl Sulf1/2 cKO (red *), or as indicated. *, **, ***, **** indicated p < 0.05, < 0.01, < 0.001, and <0.0001 respectively. **n**, hOPCs were co-infected with lentiviral SULF2 shRNAi or scrambled control and lentiviral TCF/LEF reporter virus and exposed to WNT3A (50ng/mL) or vehicle control. Two-way RM ANOVA followed by Holm-Sidak post-test (n = 4 fetal human samples, mean ± SEM normalized to control). **o**, hOPCs infected with lentiviral BMP reporter and SULF2 KD virus. SULF2 KD reduced BMP activity in hOPCs following BMP7 treatment (50 ng/ml) (n = 5, p < 0.05 two-way RM ANOVA Holm-Sidak post-test). Two-way ANOVA revealed significant main effects for ligand treatment, SULF2 KD and the interaction for both WNT and BMP. **p**, quantification of O4^+^ oligodendrocyte differentiation following SULF2 KD and BMP7 treatment (5 – 50 ng/mL BMP7) (n = 5-6 fetal human samples). SULF2 knockdown attenuated the effects of WNT/BMP signaling and significantly increased differentiation of hOPCs in the presence of BMP (Two-way RM ANOVA, Holm-Sidak p < 0.05). * and ** indicate p < 0.05 and p < 0.01, respectively. Scale: 20 μm (**a-j**).

As previously shown with genetic activation of WNT signaling (Fancy et al., 2009), pharmacological WNT activation via CHIR-99021 (3 µM) in wild-type animals resulted in significantly impaired differentiation with reduced density of CC1^+^Olig2^+^ oligodendrocytes (n = 4 and 8 respectively, Holm-Sidak multiple comparison test p < 0.05) (**Table S2**) (**Fig. 5l**) and reduced proportion of CC1^+^Olig2^+^ oligodendrocytes among total Olig2^+^ cells (Holm-Sidak p < 0.001) (**Fig. 5m**) following demyelination. Likewise, tankyrase inhibitor XAV939 (0.1 µM) which blocks WNT signaling significantly promoted oligodendrocyte differentiation in wild-type mice measured as CC1^+^Olig2^+^ density and CC1% at 7 dpl (Hold-Sidak p<0.0001), as previously shown (Fancy et al., 2011). As described above, *Sulf1/2* cKO significantly increased both metrics of differentiation (**Table S2**). In the context of *Sulf1/2* cKO, WNT activation (CHIR-99021) was unable to block oligodendrocyte differentiation (Olig2^+^CC1^+^ density p = 0.94, CC1% p = 0.89) (**Fig. 5l-m**). Likewise, additional WNT antagonism by XAV939 did not further potentiate the increase in oligodendrocyte differentiation afforded by *Sulf1/2* cKO (Olig2^+^CC1^+^ density p = 0.90, CC1% p = 0.99). Thus, interaction between WNT signaling and sulfatase was clear and strongly supported the hypothesis that *Sulf1/2* ablation acts to promote oligodendrocyte differentiation via modulation of WNT signaling.

Next, we used A01 (100 nM) to inhibit E3 ligase Smurf1 and thereby activate intracellular BMP signaling (Cao et al., 2014) and BMP type I receptor antagonist LDN193189 (100 nM) to antagonize BMP signaling (**Fig. 5a-m**). In wild-type animals, pharmacological BMP activation (A01) significantly decreased Olig2^+^CC1^+^ oligodendrocyte density and CC1^+^ percentage (Holm-Sidak p < 0.05 and p < 0.001) (**Table S2**) (**Fig. 5l-m**). In contrast, BMP receptor antagonist (LDN) increased all three parameters, total Olig2^+^ density (Holm-Sidak p < 0.001) (**Fig. 5k**), CC1^+^ density (p < 0.0001) (**Fig. 5l**) and % CC1 (p < 0.0001) (**Fig. 5m**). In *Sulf1/2* cKO mice, intracellular BMP activation via A01 treatment reduced oligodendrocyte differentiation (Olig2^+^CC1^+^ density and CC1%) compared to *Sulf1/2* cKO alone (**Table S2**) (**Fig. 5l-m**). However, this level of differentiation was significantly enriched compared to wild type indicating that *Sulf1/2* cKO effect was still effective in the context of BMP excessive activation. In contrast, *Sulf1/2* cKO did not further enhance oligodendrocyte differentiation in animals treated with BMP receptor antagonist (LDN) (Hold Sidak, Olig2^+^CC1^+^ density p = 0.99, CC1% p = 0.18) (**Fig. 5l-m**). The interaction between BMP receptor antagonist treatment and sulfatase suggests that sulfatase acts via BMP signaling.

As modulation of these pathways could occur at multiple levels, we next asked whether sulfatase modulated WNT and BMP signaling directly in human OPCs. To this end, scrambled control or SULF2 knockdown (KD) hOPCs were transduced with viral reporters for WNT and BMP signaling (see methods). WNT-dependent TCF reporter luciferase was >11-fold induced following WNT3a treatment (n = 4 fetal samples) (**Fig. 5n**). Strikingly, SULF2 KD significantly attenuated WNT induction by more than 40%. Likewise, BMP response element (BRE)-dependent luciferase was increased >18-fold by BMP7 treatment and this BMP-induced BRE luciferase activity was significantly inhibited by SULF2 KD (**Fig. 5o**). Two-way ANOVA revealed a significant interaction effect between SULF2 KD and both WNT and BMP signaling (p < 0.05) and was consistent with a direct effect of sulfatase on these signaling pathways. To further test whether SULF2 KD could directly influence the effects of BMP signaling on hOPC fate, we treated SULF2 KD and scrambled infected hOPCs with BMP7 and assessed the effects on O4^+^ oligodendrocyte differentiation (n = 5-6 fetal samples) (**Fig. S6**). As shown previously (Sim et al., 2011), BMP7 treatment inhibited O4^+^ differentiation from hOPCs (Two-way ANOVA, p < 0.001) (**Fig. 5p**). Importantly, SULF2 KD significantly rescued the effects of BMP7 (Hold-Sidak test, p < 0.01). As such, sulfatase expressed by hOPCs directly potentiates both WNT and BMP signaling and is consistent with sulfatases exerting cell-autonomous role in OPCs.

### PI-88 modulates OPCs sulfation and regulates BMP and WNT signaling in hOPCs

As sulfatases act in the extracellular space, they represent a pharmacologically relevant target for manipulation in demyelinating disease. PI-88, a highly sulfated structural mimetic of heparan sulfate acts as a non-cleavable substrate and inhibitor for sulfatases (Parish et al., 1999, Yu et al., 2002) and is currently in clinical trials for cancer therapy. We first assessed basal HS sulfation on rat CG4 and human primary OPCs by flow cytometry using RB4CD12 antibody that recognizes highly sulfated HS GAG motifs (Jenniskens et al., 2000) (**Fig. 6a, b**). Treatment with PI-88 caused a progressive and persistent enrichment of HS sulfation at 30-min and 24 hours (**Fig. 6b**).

**Figure 6.**
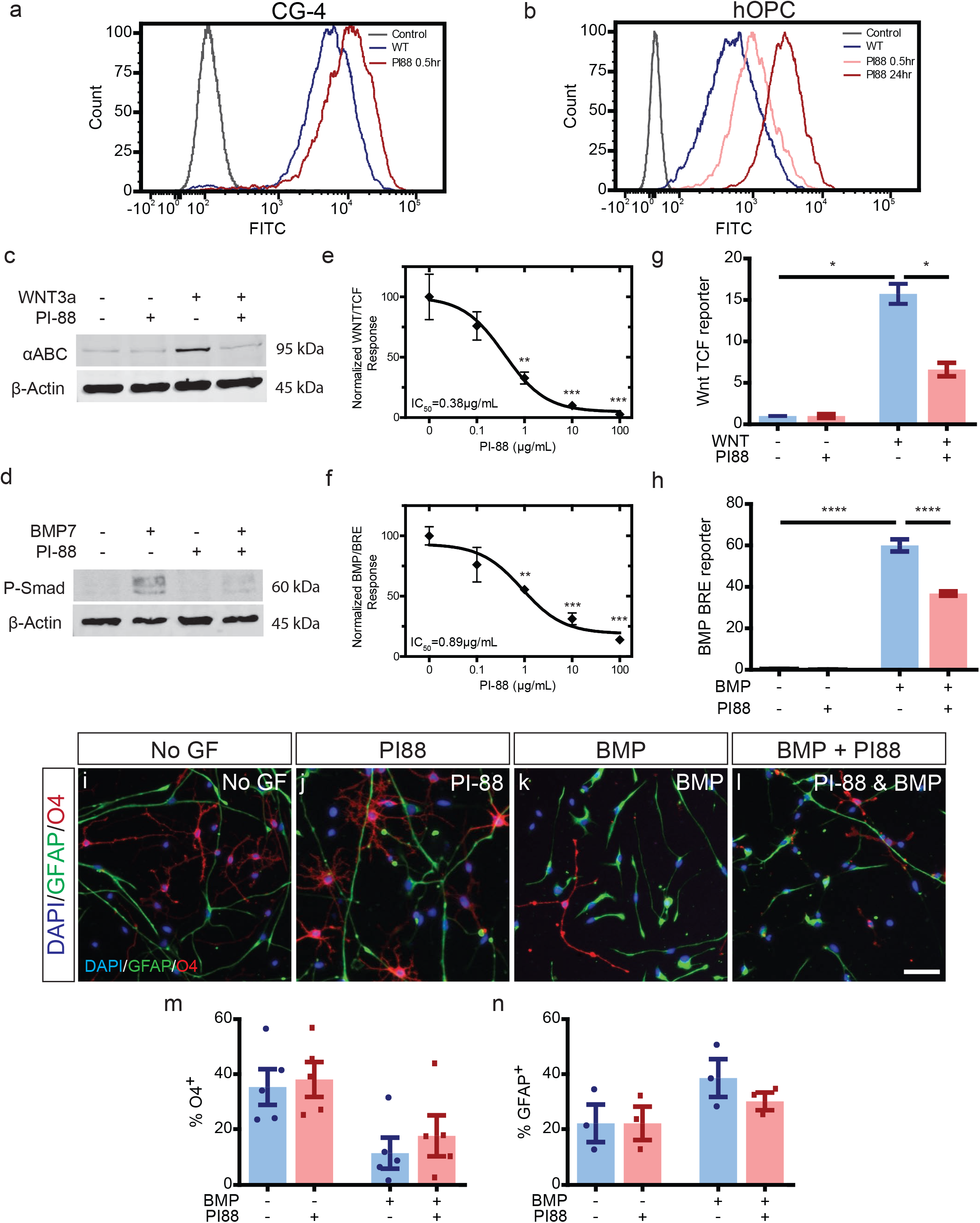
PI-88 inhibits sulfatases leading to increased HSPG sulfation, and inhibition of WNT and BMP signaling in OPCs. **a-b**, Flow cytometry of HS sulfation using RB4CD12 antibody on rat CG-4 (**a**) and human primary (**b**) OPCs. No antibody control shown (green). PI-88 treatment (100 μg/ml) resulted in a clear increase in OPC sulfation in a time-dependent manner. **c-d**, Western blots for active β-catenin (αABC) and phosphorylated Smad 1/5 (P-Smad) following treatment of rat CG-4 OPCs treated with BMP7, WNT3a (both 50ng/mL) and/or PI-88 (100 μg/ml). **e-f**, Dose-response curves for WNT and BMP reporter activity in the presence/absence of PI-88 (20 hours following treatment, n = 3, one-way ANOVA). ** and *** indicate significant effect of PI-88 by Dunnett’s post-test p < 0.01, < 0.001. **g**, human primary OPCs were infected with lentiviral TCF reporter and treated with PI-88 (2 µg/mL) and/or WNT3a (50 ng/ml) (n=3 fetal human samples, normalized mean ± SEM). **h**, similarly, hOPCs were infected with BMP response element dependent reporter and treated with PI-88 and/or BMP7 (n = 3). Two-way ANOVA revealed significant effects of WNT/BMP and PI-88 treatment, *, *** indicates Holm-Sidak post-test p < 0.05, p < 0.001, respectively. **i-n**, the effect of PI-88 treatment on hOPC differentiation was assessed in the context of inhibitory BMP treatment. hOPC were treated with BMP and/or PI-88 and O4^+^ oligodendrocyte (red) and GFAP^+^ astrocyte (green) differentiation assessed after removal of mitogens. Quantification of O4% (**m**, n = 5 fetal samples) and GFAP% (**n**, n = 3), mean ± SEM shown. Two-way RM ANOVA revealed significant effects of BMP and PI-88 on both O4 and GFAP% (main effect p < 0.05). Scale: 50 µm.

We next examined whether PI-88 via induced increased HS sulfation could inhibit WNT and BMP signaling in OPCs. In rat CG4 cells, activated β-catenin, a downstream effector of canonical WNT signaling was substantially induced following WNT3a treatment (**Fig. 6c**). Importantly, PI-88 treatment effectively blocked the effect of WNT3a. Likewise, PI-88 attenuated BMP7-induced Smad 1/5 phosphorylation (pSmad) (**Fig. 6d**). Using luciferase-expressing lentiviral reporters of WNT and BMP signaling, we found PI-88 exhibited a clear dose-dependent effect and antagonized WNT and BMP signaling with IC50s of 0.38 μg/ml and 0.89 μg/ml for WNT3a and BMP7 respectively (**Fig. 6e-f**). Interestingly, when we compared the effect of PI-88 treatment on low and high dose WNT/BMP-induced luciferase we observed a downward shift of the dose-response curve following PI-88 treatment (data not shown). This suggests that the effect of PI-88 is noncompetitive and is consistent with inhibition of sulfatase rather than direct agonist binding or another receptor-mediated mechanism.

Next, we explored the effects of PI-88 on human primary OPC signaling. Lentiviral reporter infected hOPCs were treated with BMP/WNT ligands and/or PI-88. BMP and WNT ligand treatment (50 ng/ml) induced robust >15-fold increase in reporter activity (two-way ANOVA, p < 0.05) that was significantly attenuated by PI-88 (**Fig. 6g-h**). PI-88 treatment alone did not alter BMP or WNT-dependent luciferase activity. Thus, we concluded that PI-88 was capable of inhibiting BMP and WNT signaling in both rat and hOPCs.

### PI-88 rescues the BMP7 mediated inhibition of hOPC differentiation *in vitro*

The inhibitory effects of PI-88 treatment on BMP-induced transcription suggested that PI-88-mediated inhibition of sulfatase activity could prevent BMP-mediated effects on OPC cell fate (Sim et al., 2011). hOPCs were cultured following mitogen withdrawal to induce oligodendrocyte differentiation. As described above, BMP7 (5 ng/ml) blocked O4^+^ differentiation (n = 4 fetal samples, two-way RM ANOVA p < 0.001) (**Fig. 6m**). Following BMP7 treatment, PI-88 treatment significantly increased O4^+^ differentiation (Holm-Sidak p<0.05). Likewise, BMP7 induced GFAP^+^ astrocyte fate from hOPCs (**Fig. 6n)**. Following PI-88 treatment, BMP7 was unable to significantly increase astrocyte differentiation. In the absence of BMP7, PI-88 did not significantly alter hOPC fate. Taken together, these results suggest that pharmacologically biasing hOPCs toward a state of elevated 6-O sulfation is phenotypically relevant, opposing the BMP7-mediated induction of astrocytic fate *in vitro*.

### PI-88 promotes oligodendrocyte differentiation and remyelination following focal demyelination

As PI-88 was able to effectively modulate HS sulfation and antagonize WNT and BMP signaling *in vitro*, we next asked whether sulfatase inhibition by PI-88 could modulate remyelination. PI-88 (10 µg/ml) or saline was co-injected with lysolecithin. At 3 dpl, PI-88 did not influence lesion size, microglial infiltration or astrogliosis. Loss of oligodendrocytes and OPCs within the lesion was likewise complete in both groups (**Fig. S7**). As such lesion formation was indistinguishable between PI-88 treated and untreated animals. At 5 dpl, during OPC recruitment, we observed a significant >25% increase in Olig2^+^ density following PI-88 treatment (n = 6-7 animals, t-test p < 0.01) (**Fig. 7a-b, k**) that was not due to increased proliferation (EdU%, p = 0.78) (**Fig. 7l**). Similar to our observations following *Sulf1/2* cKO, PI-88 treatment did not notably influence astrocytic or microglial responses at 7 dpl (**Fig. 7g-j and Fig. S8**). At 7 dpl, the density of oligodendrocytes, defined as the density of either *Plp1*^+^ or CC1^+^Olig2^+^ cells, was significantly increased by PI-88 (n = 4, p < 0.05) (**Fig. 7m-n**). Similar to *Sulf1/2* cKO, the overall Olig2^+^ oligodendrocyte lineage cell density was significantly increased by PI-88 treatment (p < 0.05) (**Fig. 7n**). Consistent with accelerated differentiation, the percentage of CC1^+^ oligodendrocytes among total Olig2^+^ was significantly increased by PI-88 (n = 6-7) (**Fig. 7o**). Thus, PI-88 treatment acts in a similar manner to *Sulf1/2* cKO resulting in increased oligodendrocyte density following demyelination.

**Figure 7.**
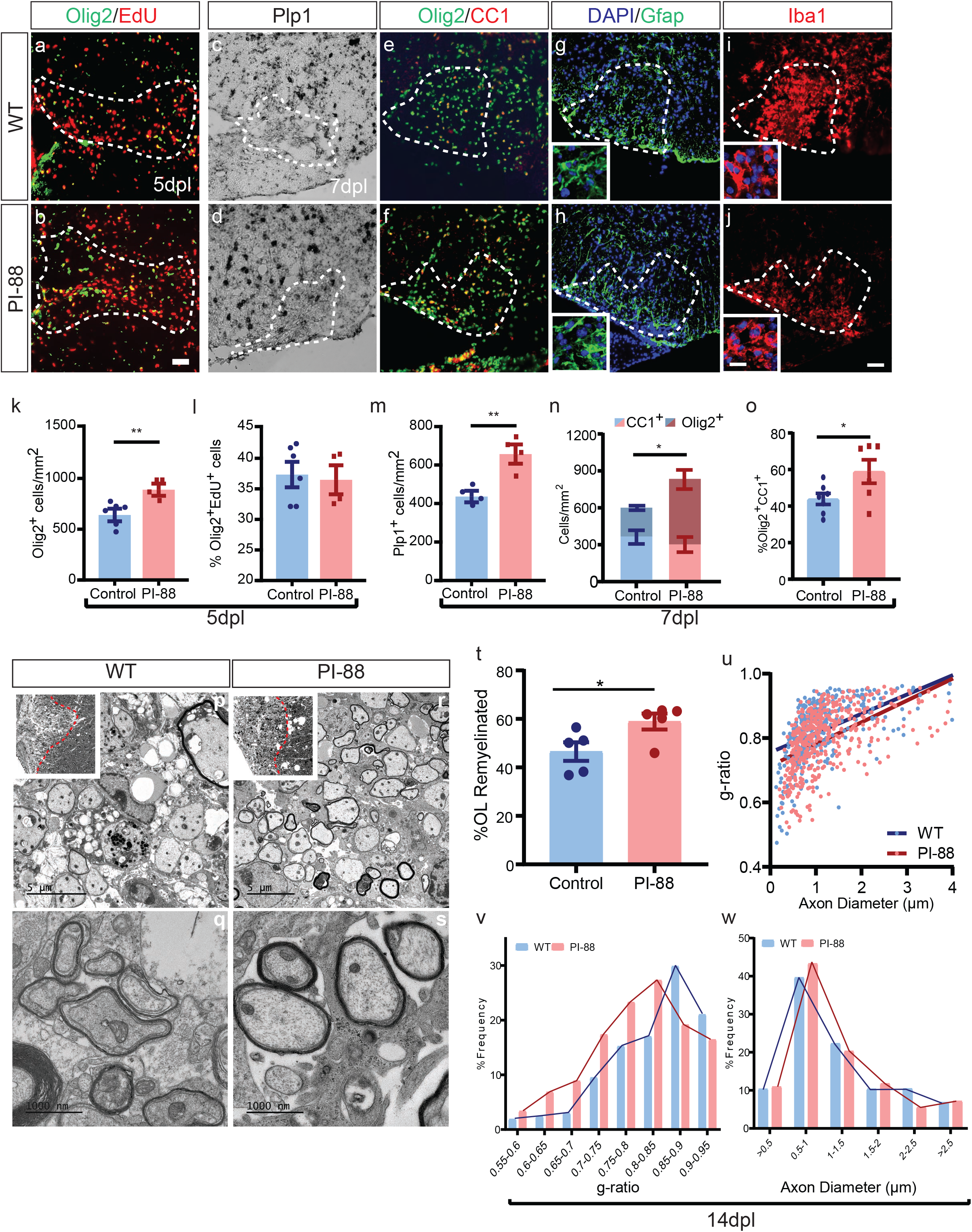
Sulfatase inhibitor PI-88 promotes OPC differentiation and remyelination following demyelination. PI-88 was injected into demyelinated lesions in young adult mouse spinal cord at the time of lysolecithin injection. **a-b**, Olig2^+^ recruitment and proliferation (EdU^+^Olig2^+^) was assess at 5 days post-lesion (dpl) by widefield epifluorescence. **c-j**, oligodendrocyte lineage density, differentiation and the microglial/astrocyte response was assessed at 7 dpl. Oligodendrocyte density and differentiation was assessed by *Plp1 in situ* (**c-d**), and Olig2/CC1 immunofluorescence (**e-f**). **g-j**, Astrocyte and microglial responses were assessed by Gfap and Iba1, respectively and image inserts shows higher magnification image to show morphology. **k-l**, quantification of Olig2^+^ density and percentage of EdU^+^ cells among total Olig2 at 5 dpl (n = 6-7 animals, mean ± SEM). *, ** indicate t-test p < 0.05 and 0.01, respectively. **m-o**, at 7 dpl quantification of *Plp1*^*+*^ cell density (**m**), Olig2^+^ and CC1^+^Olig2^+^ cell density (**n**), and percentage of CC1^+^ cells within the Olig2 population (**o**) (n = 4-6 animals). **p-w**, Analysis of remyelination by EM at 14 dpl (n = 5 animals/group). **p**,**r**, Semithins inserts showing lesions in ventrolateral white matter. **t**, Proportion of remyelinated axons (* indicates t-test p < 0.05), **u**, relationship between axon diameter and *g*-ratio (linear regression shown). Frequency distribution of axonal *g*-ratio (**v**) and axon diameter (**w**) in lesion. Scale: 20 μm (**a-j**) and 5 μm (**p-r**) and 1000 nm(**q-s**).

Next, we examined whether enhanced differentiation following PI-88 treatment resulted in accelerated remyelination (**Fig. 7p-s**). We found that PI-88 treatment significantly increased the proportion of remyelinated axons at 14 dpl (n = 5 animals per group, t-test p < 0.05) (**Fig. 7t**). Linear regression analysis of axon diameter vs. *g*-ratios demonstrated a significant reduction of g-ratio following PI-88 treatment (**Fig. 7u**). Likewise, the distribution of *g*-ratios binned by axon diameter was left shifted and indicated increased myelin sheath thickness in PI-88 treated mice (**Fig 7v**). Similar to *Sulf1/2* cKO, PI-88 treatment did not induce axonal swelling confirmed by axon diameter frequency distribution (**Fig 7w**). Together, these data demonstrate that PI-88 treatment promoted both OPC differentiation and accelerated remyelination following demyelination.

### PI-88 induced oligodendrocyte differentiation following differentiation is mediated via inhibition of sulfatase

To determine the mechanism by which PI-88 increased oligodendrocyte differentiation, and whether the effects of PI-88 were dependent on OPC-expressed sulfatase, we treated mice with PI-88 in the context of OPC-specific sulfatase deletion (i.e. *Sulf1/2* cKO). We induced lysolecithin lesions in *Sulf1/2* cKO and treated with PI-88 (n = 4-5 animals per group) (**Fig. 8**). As described above, PI-88 treatment or *Sulf1/2* cKO alone significantly increased Olig2^+^ oligodendrocyte lineage density (**Fig. 8m**), Olig2^+^CC1^+^ density (**Fig. 8n**), and the proportion of CC1^+^ oligodendrocytes (**Fig. 8n**). Importantly, we did not observe an additive effect of PI-88 treatment on *Sulf1/2* cKO mice for these parameters (**Fig. 8m-o**). Two-way ANOVA revealed a highly significant interaction between genotype and PI-88 treatment on all parameters (**Table S4**). Thus, PI-88 likely lead to increased oligodendrocyte differentiation primarily via inhibition of OPC-expressed *Sulf1/2*.

**Figure 8.**
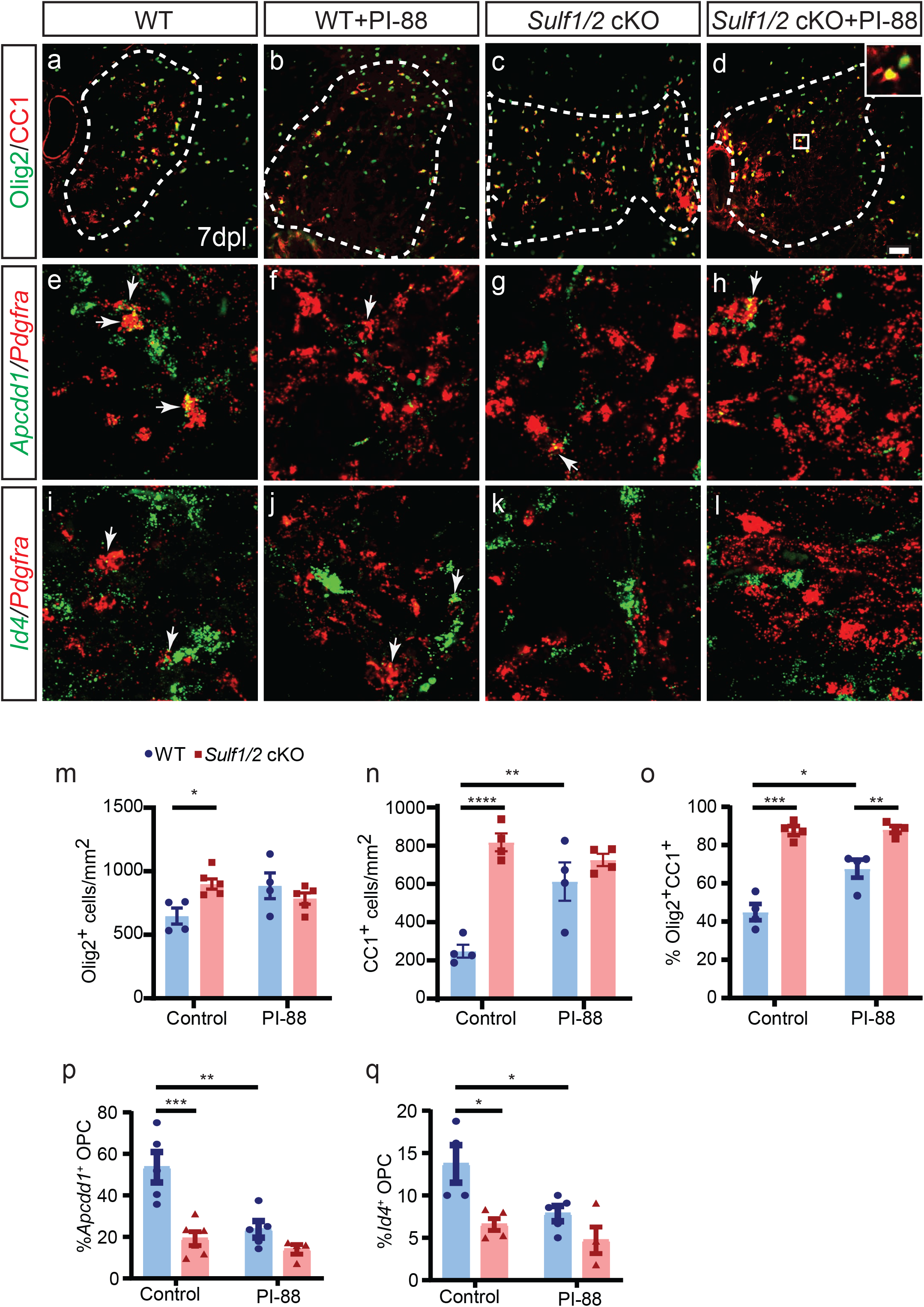
PI-88 treatment accelerated OL differentiation via inhibition of *Sulf1/2* following demyelination. WT and *Sulf1/2* cKO mice underwent focal spinal cord demyelination with or without PI-88 injection and were sacrificed at 7 days post-lesion. **a-d**, Olig2 (green)/CC1 (red) immunofluorescence. **e-l**, Confocal RNAscope *in situ* hybridization for OPC-expressed *Pdgfra* (red) and either WNT target gene *Apcdd1* (green, **e-h**) or BMP target gene *Id4* (green, **i-l**). Arrows indicate double labelled OPCs. **m-o**, Quantification of oligodendrocyte lineage density (Olig2^+^, **m**), post-mitotic CC1^+^Olig2^+^ cells (**n**), and percentage of CC1^+^ cells within the Olig2 population (**o**) (n = 4-5 animals per group). Quantification of WNT pathway activity *Appcd1*% (**p**) and *Id4*% (**q**) among *Pdgfra*^*+*^ OPCs (n = 4-6 animals). Mean ± SEM shown. *, **, ***, **** indicate Two-way ANOVA Holm-Sidak post-test p < 0.05, <0.01, <0.001, and < 0.0001 respectively. Scale: 20 µm.

Finally, to determine if PI-88 might similarly regulate WNT and BMP signaling and exert a pro-differentiative effect via WNT/BMP antagonism, we examined the expression of WNT and BMP target genes following *Sulf1/2* cKO and/or PI-88 treatment using *in situ* hybridization (**Fig. 8e-l**). Similar to our results using pharmacological manipulation of WNT and BMP pathways, we observed significantly decreased expression of *Apcdd1* (WNT target) and *Id4* (BMP target) in *Pdgfra*^*+*^ OPCs in *Sulf1/2* cKO mice (two-way ANOVA, *Sulf1/2* cKO main effect p < 0.01, n = 4-6 animals per group) (**Fig 8p-q**). Likewise, PI-88 treatment alone significantly inhibited expression of both genes in *Pdgfra*^*+*^ OPCs (two-way ANOVA, PI-88 effect p < 0.05). Combined treatment of PI-88 in *Sulf1/2* cKO mice did not significantly decrease WNT or BMP target gene expression any further compared to *Sulf1/2* cKO alone (Holm-Sidak, p > 0.5). These observations suggest that PI-88 acts via OPC-expressed *Sulf1/2* to influence BMP and WNT signaling in OPCs and thereby promote myelin repair.

## DISCUSSION

The inhibitory tissue environment of chronic demyelinated lesions acts to prevent efficient myelin repair and remyelination in MS (Lau et al., 2013). The cellular environment and local extracellular matrix determine the availability and signaling of multiple growth factor and cytokine pathways. Here, we show that sulfation of heparan sulfate proteoglycans (HSPG) critically influences the paracrine signaling environment surrounding OPCs and can be modulated to improve recruitment and formation of new oligodendrocytes leading to accelerated myelin repair. These data demonstrate that extracellular sulfatases expressed by human and mouse OPCs promote inhibitory WNT and BMP signaling following demyelination and that ablation of sulfatase function either by conditional genetic deletion or pharmacological inhibition can enhance remyelination.

The mammalian genome contains two sulfatase genes which share substrate specificity but differ in expression pattern in a cell- and tissue-specific manner (Ai et al., 2006, Morimoto-Tomita et al., 2002). We found that OPCs express high levels of sulfatases that are eliminated during oligodendrocyte differentiation. During mouse CNS development, both sulfatases were expressed by OPCs and immature oligodendrocytes. In contrast, human OPCs express SULF2 more than 10^4^-fold higher than SULF1. The significance of this species-difference in expression is unclear but has significant implications for the design of sulfatase-specific small molecules. In contrast, the similarities in terms of resultant phenotype following sulfatase deletion suggest an important conservation of function between human and mouse signaling. In the adult mouse CNS, Sulf1/2 expression is largely restricted to OPCs and a subpopulation of cortical layer V neurons, as previously described (Zeisel et al., 2015). Neuronal Sulf2 expression suggests that the heparanome within cortical demyelinated lesions differs from subcortical white matter and may potentiate inhibitory signals that contribute to failed remyelination. Intriguingly, sulfatase expression remained predominantly restricted to OPCs following demyelination in both mouse and human lesions. While the specific HSPG core proteins that undergo sulfatase-mediated editing are unknown, syndecan-3 (SDC3) is noteworthy due to its high expression in fetal and adult human OPCs (Sim et al., 2009, Sim et al., 2011) and in cultured murine cells (Winkler et al., 2002).

The role of heparanome sulfation has not been previously studied in the context of demyelination. Constitutive sulfatase expression by OPCs suggests these enzymes prevent complete 6-O sulfation and our functional studies indicate that increased HSPG sulfation is associated with inhibition of various extracellular signaling cascades. The resting state of OPC heparanome is predominately composed of highly sulfated HexA2S-GlcNS6S trisulfated disaccharide units which are specific among glial subtypes (Stringer et al., 1999). As such, sulfatases are ideally situated to modulate OPC signaling. The generation of highly sulfated HSPGs is catalyzed by sulfotransferases. These enzymes, expressed at high levels by OPCs, are localized to the Golgi apparatus and do not act once HSPGs are presented on the cell surface (Habuchi et al., 2004). As such, the increased prevalence of surface HS sulfation following sulfatase inhibition is most likely due to *de novo* HSPG presentation and suggests a rapid turnover of OPC-expressed HSPGs.

In the adult CNS, sulfatase deletion in OPCs results in a substantial increase in recruitment and differentiation of oligodendrocyte lineage cells following demyelination leading to improved remyelination. Interestingly, antisense Sulf1 treatment during embryonic development results in reduced migration (Kakinuma et al., 2004). Similarly, we observe that SULF KD in hOPCs results in perturbed migration in response to PDGF-AA (data not shown). As HSPG sulfation is expected to influence multiple pathways, we observed distinct effects on OPC dynamics following demyelination. In addition to accelerated differentiation (assessed by density and proportion of post-mitotic oligodendrocytes), we noted that sulfatase deletion resulted in progressive recruitment of Olig2^+^ cells such that the density of oligodendrocyte lineage cells progressively increased from 5-14 days post lesion. This contrasts with wild-type lesions which typically reach maximal density at 5 days. The mechanisms by which overall recruitment of OPC and subsequent generation of oligodendrocytes remain unclear and are likely complex due to the number of signaling pathways that may be influenced by sulfatase activity. Regardless, the effect of dual sulfatase deletion on oligodendrocyte number was striking as we observed a more 2-fold increase in oligodendrocyte number. Consistent with shared substrate specificity, the effect of individual sulfatase deletion was consistent with a model in which sulfatases act in a dose-dependent manner. As such, once a lower limit of sulfatase activity was reached, oligodendrocyte recruitment and differentiation were increased. The dose-dependent effects of sulfatase inhibition *in vitro* is consistent with this model and supports the potential for future therapeutic intervention. Importantly, the effect of conditional deletion of *Sulf1/2* in NG2-expressing cells was restricted to OPCs as we did not observe pericyte Sulf1/2 expression either in normal CNS or following demyelination with similar results observed in purified human OPCs. As pericytes did not express Sulf1/2, conditional deletion of sulfatase in NG2-expressing cells resulted in specific deletion of OPC-expressed sulfatase activity.

HSPG sulfation is known to regulate several signaling pathways (Rosen and Lemjabbar-Alaoui, 2010). Here, we show both WNT and BMP pathways are inhibited by sulfatase inhibition in OPCs. Canonical WNT signaling prevents efficient oligodendrocyte differentiation following demyelination (Fancy et al., 2009, Fancy et al., 2011). WNT target genes are activated in OPCs following human white matter injury and correlate with failed differentiation and repair (Fancy et al., 2014). We found that endogenous sulfatases promote WNT signaling and sulfatase inhibition decreases WNT target gene expression in OPCs. This likely occurs in a ligand-receptor dependent manner as TCF/LEF transcriptional activity was dependent on SULF2 expression in purified human OPCs.

We observed a highly significant interaction between sulfatase function and pharmacological manipulation of WNT signaling. CHIR-99021 acts as a GSK-3β inhibitor to potentiate canonical WNT signaling. As *Sulf1/2* cKO was observed to completely block the effect of the GSK-3β antagonist, these data suggest that canonical WNT signaling acts to inhibit oligodendrocyte differentiation and remyelination via secondary pathways which are themselves dependent on and modulated by *Sulf1/2*. In purified rat OPCs, the inhibitory effect of WNT3a treatment on differentiation occurs in a BMP-dependent manner (Feigenson et al., 2011). In contrast, recombinant human WNT3a treatment of purified hOPCs while effectively activating TCF/LEF transcription did not alter basal oligodendrocyte differentiation (data not shown) suggesting the need for engagement of additional pathways to elicit the inhibitory WNT effect. Together with our data, these observations suggest that WNT may be acting via and dependent on BMP and other extracellular signaling following demyelination. In addition to these paracrine mechanisms, it is likely that sulfatases also directly modulate WNT signaling. Supporting a direct role of sulfatase manipulation on WNT signaling in demyelination, we observed the lack of an additive effect of XAV939 treatment and sulfatase deletion. In addition, we found that WNT target gene expression is reduced following *Sulf1/2* cKO and that WNT-dependent transcriptional activity in hOPC was similarly reduced following SULF2 KD. While WNT pathways is therefore likely a key target of sulfatases following demyelination, WNT antagonist treatment did not entirely phenocopy *Sulf1/2* cKO, as treatment with XAV939 did not increase overall Olig2 density whereas *Sulf1/2* cKO resulted in a significantly increased density.

Our data support a clear role for sulfatase mediated potentiation of BMP signaling following demyelination. Genetic sulfatase deletion in OPCs resulted in impaired BMP-response element-dependent transcription and reduced Id4 expression by OPCs *in vivo*. BMP signaling is up regulated following demyelination (Ara et al., 2008, Sabo et al., 2011) and is active in MS lesions (Deininger et al., 1995). We found that BMP receptor antagonist increased both Olig2 recruitment and oligodendrocyte differentiation, as shown (Govier-Cole et al., 2019). Sulfatase deletion in the context of BMP blockade did not further increase differentiation suggesting this process is dependent on BMP signaling. In contrast, intracellular activation of BMP signaling by treatment with A01, a Smurf1 E3 ligase specific antagonist, could not be effectively compensated for by sulfatase deletion. Given the intracellular mode of action for A01, this is entirely consistent with a model in which HSPG sulfation regulations BMP ligand/receptor accessibility.

In addition to WNT and BMP, modulation of HSPG sulfation by sulfatases regulates several additional signaling cascades that influence the demyelinated lesion microenvironment. Previously, Sulf activity has been to shown to potentiate PDGFαR signaling in glioblastoma (Phillips et al., 2012). PDGF is the principle OPC mitogen during development (Sim et al., 2011, Sim et al., 2006, Noble et al., 1988, Raff et al., 1988) but does not appear to be rate-limiting following murine spinal cord demyelination (Calver et al., 1998, Woodruff et al., 2004). In addition to PDGF, previous studies suggest that sulfatase activity inhibits FGF2 signaling (Otsuki et al., 2010, Wang et al., 2004, Lamanna et al., 2006, Holst et al., 2007, Seffouh et al., 2013) and pharmacologically reduced sulfation blocks FGF responsiveness in OPCs (Bansal et al., 1996, Fortin et al., 2005). As FGFR ablation in oligodendrocyte lineage cells results in hypomyelination (Furusho et al., 2012), inhibition of sulfatase leading to increased FGF signaling is consistent with our observed acceleration of remyelination and increased myelin sheath thickness following *Sulf1/2* cKO. In addition to PDGF and FGF2, heparan sulfatase regulation of inflammatory signaling cascades such as IFN-γ have been described (Lortat-Jacob et al., 1991). As such, HSPG 6-O sulfation via regulation by OPC-expressed sulfatases provide the basis for the coordinated regulation of the lesion microenvironment.

Given that sulfatases are secreted into the extracellular milieu, it is likely that paracrine effects of sulfatase inhibition may influence other glial and inflammatory cell signaling following demyelination. Although a broad assessment of microglial number via Iba1 immunoreactivity and astrogliosis via Gfap did not indicate a major effect following sulfatase deletion, we cannot exclude the possibility that paracrine effects may influence the infiltration, proliferation, or activation of microglia and other immune cells within the lesion environment. Importantly, while lysolecithin-induced demyelination provides the ideal model to assess the effects of sulfatase deletion on the cellular dynamics of remyelination, paracrine influences of sulfatase inhibition cannot be ruled out due to the relative absence of adaptive immune system related signaling in this model.

PI-88 is a heparin mimetic acting as a competitive antagonist to sulfatases (Parish et al., 1999). The effects of PI-88 treatment on OPC dynamics following lysolecithin demyelination essentially phenocopied that of OPC-specific sulfatase deletion. Importantly, when PI-88 treatment was combined with *Sulf1/2* cKO, there were no additive effects and the PI-88 did not further influence OPC dynamics or differentiation. This is consistent with the effects of PI-88 being mediated solely via inhibition of OPC-expressed sulfatase. However, we cannot exclude the possibility that heparanase antagonism by PI-88 could influence the lesion environment (Karoli et al., 2005).

By modulating the local OPC niche in the context of demyelination, we have found a single target capable of modulating several signaling cascades in concert. Previous remyelination approaches have targeted single signaling pathways such as LINGO-1 (Mi et al., 2007), WNT (Fancy et al., 2011), RXRγ, muscarinic receptor antagonists (Mei et al., 2014, Deshmukh et al., 2013, Abiraman et al., 2015), amongst others. Other approaches such as miconazole or clobetasol have less defined mechanisms of action (Najm et al., 2015). While these and other small molecules have been discovered that promote OPC differentiation and remyelination in rodent models, given the apparent diversity of inhibitory cascades involved, it seems unlikely that targeting single pathways will yield an effective clinical intervention. As such, PI-88 may allow simultaneous modulation of several distinct signaling inputs in favor of a single biological goal, a unique approach to the treatment of demyelinating disease. In addition as cell therapies for myelin disorders approach clinical translation (Franklin and Goldman, 2015), modulation of the heparanome might represent a viable approach to overcome the local inhibitory environment into which cells are engrafted. Future studies will need to directly address the clinical utility of PI-88, and whether PI-88 or another Sulf-targeting compound could function synergistically with one or more of the previously described compounds to exact efficient myelin repair. Systemic PI-88 (Muparfostat) is currently being investigated in various oncology-related clinical trials (clinicaltrials.gov), with a favorable safety profile reported (Chen et al., 2017).

Taken together, OPC-expressed sulfatases by regulating their local heparanome, potentiate inhibitory signals present in demyelinating lesions that prevent efficient myelin repair. Treatment with PI-88 represented a potent means to inhibit sulfatases and support the potential for therapeutic approaches aimed at sulfatase inhibition. CNS optimized delivery of PI-88 or similar heparan mimetics may represent a potent and multifaceted approach to achieve enhancement of endogenous myelin repair in diseases such as multiple sclerosis.

## Acknowledgements

This work was supported by NINDS grant (R01NS104021), NCATS (UL1TR001412), National Multiple Sclerosis Society (RG 5505-A-2, RG-1701-26750), the Kalec Multiple Sclerosis Foundation, the Change MS Foundation, the Skarlow Memorial Trust, and the Empire State Stem Cell Fund through New York State Department of Health Contract (C028108). JJP (Polanco) received additional support from NIGMS (R25GM09545902), NCATS (UL1TR001412-S1), and New York State Department of Health (Empire State Stem Cell Fund) through the Stem Cells in Regenerative Medicine Fellowship (NYSTEM C30290GG). JJP (Phillips) received support from NCI (1U01CA229345). We acknowledge the assistance of the Confocal Microscope and Flow Cytometry Facility in the School of Medicine and Biomedical Sciences, University at Buffalo.

## REFERENCES

Abiraman, K., Pol, S. U., O’Bara, M. A., Chen, G. D., Khaku, Z. M., Wang, J., Thorn, D., Vedia, B. H., Ekwegbalu, E. C., Li, J. X., Salvi, R. J. & Sim, F. J. 2015. Anti-muscarinic adjunct therapy accelerates functional human oligodendrocyte repair. J Neurosci, 35, 3676–88.

Ai, X., Do, A. T., Kusche-Gullberg, M., Lindahl, U., Lu, K. & Emerson, C. P., JR. 2006. Substrate specificity and domain functions of extracellular heparan sulfate 6-O-endosulfatases, QSulf1 and QSulf2. J Biol Chem, 281, 4969–76.

Ai, X., Kitazawa, T., Do, A. T., Kusche-Gullberg, M., Labosky, P. A. & Emerson, C. P., JR. 2007. SULF1 and SULF2 regulate heparan sulfate-mediated GDNF signaling for esophageal innervation. Development, 134, 3327–38.

Ara, J., See, J., Mamontov, P., Hahn, A., Bannerman, P., Pleasure, D. & Grinspan, J. B. 2008. Bone morphogenetic proteins 4, 6, and 7 are up-regulated in mouse spinal cord during experimental autoimmune encephalomyelitis. J Neurosci Res, 86, 125–35.

Bansal, R., Kumar, M., Murray, K., Morrison, R. S. & Pfeiffer, S. E. 1996. Regulation of FGF receptors in the oligodendrocyte lineage. Mol Cell Neurosci, 7, 263–75.

Biechele, T. L. & Moon, R. T. 2008. Assaying beta-catenin/TCF transcription with beta-catenin/TCF transcription-based reporter constructs. Methods Mol Biol, 468, 99–110.

Calver, A. R., Hall, A. C., Yu, W. P., Walsh, F. S., Heath, J. K., Betsholtz, C. & Richardson, W. D. 1998. Oligodendrocyte population dynamics and the role of PDGF in vivo. Neuron, 20, 869–82.

Cao, Y., Wang, C., Zhang, X., Xing, G., Lu, K., Gu, Y., He, F. & Zhang, L. 2014. Selective small molecule compounds increase BMP-2 responsiveness by inhibiting Smurf1-mediated Smad1/5 degradation. Sci Rep, 4, 4965.

Chen, P.-J., Lee, P.-H., Han, K.-H., Fan, J., Cheung, T. T., Hu, R.-H., Paik, S. W., Lee, W.-C., Chau, G.-Y., Jeng, L.-B., Wang, H. J., Choi, J. Y., Chen, C.-L., Cho, M., Ho, M.-C., Wu, C.-C., Lee, K. S., Mao, Y., Hu, F.-C. & Lai, K.-L. 2017. 624PDA phase III trial of muparfostat (PI-88) as adjuvant therapy in patients with hepatitis virus related hepatocellular carcinoma (HV-HCC) after resection. Annals of Oncology, 28.

Conway, G. D., O’Bara, M. A., Vedia, B. H., Pol, S. U. & Sim, F. J. 2012. Histone deacetylase activity is required for human oligodendrocyte progenitor differentiation. Glia, 60, 1944–53.

Deininger, M., Meyermann, R. & Schluesener, H. 1995. Detection of two transforming growth factor-beta-related morphogens, bone morphogenetic proteins-4 and -5, in RNA of multiple sclerosis and Creutzfeldt-Jakob disease lesions. Acta Neuropathol, 90, 76–9.

Deshmukh, V. A., Tardif, V., Lyssiotis, C. A., Green, C. C., Kerman, B., Kim, H. J., Padmanabhan, K., Swoboda, J. G., Ahmad, I., Kondo, T., Gage, F. H., Theofilopoulos, A. N., Lawson, B. R., Schultz, P. G. & Lairson, L. L. 2013. A regenerative approach to the treatment of multiple sclerosis. Nature, 502, 327–332.

Dillenburg, A., Ireland, G., Holloway, R. K., Davies, C. L., Evans, F. L., Swire, M., Bechler, M. E., Soong, D., Yuen, T. J., Su, G. H., Becher, J. C., Smith, C., Williams, A. & Miron, V. E. 2018. Activin receptors regulate the oligodendrocyte lineage in health and disease. Acta Neuropathol, 135, 887–906.

Fancy, S. P., Baranzini, S. E., Zhao, C., Yuk, D. I., Irvine, K. A., Kaing, S., Sanai, N., Franklin, R. J. & Rowitch, D. H. 2009. Dysregulation of the Wnt pathway inhibits timely myelination and remyelination in the mammalian CNS. Genes Dev, 23, 1571–85.

Fancy, S. P., Harrington, E. P., Baranzini, S. E., Silbereis, J. C., Shiow, L. R., Yuen, T. J., Huang, E. J., Lomvardas, S. & Rowitch, D. H. 2014. Parallel states of pathological Wnt signaling in neonatal brain injury and colon cancer. Nat Neurosci, 17, 506–12.

Fancy, S. P., Harrington, E. P., Yuen, T. J., Silbereis, J. C., Zhao, C., Baranzini, S. E., Bruce, C. C., Otero, J. J., Huang, E. J., Nusse, R., Franklin, R. J. & Rowitch, D. H. 2011. Axin2 as regulatory and therapeutic target in newborn brain injury and remyelination. Nat Neurosci, 14, 1009–16.

Feigenson, K., Reid, M., See, J., Crenshaw, I. E. & Grinspan, J. B. 2011. Canonical Wnt signalling requires the BMP pathway to inhibit oligodendrocyte maturation. ASN Neuro, 3, e00061.

Fortin, D., Rom, E., Sun, H., Yayon, A. & Bansal, R. 2005. Distinct fibroblast growth factor (FGF)/FGF receptor signaling pairs initiate diverse cellular responses in the oligodendrocyte lineage. J Neurosci, 25, 7470–9.

Franklin, R. J. 2015. Regenerative Medicines for Remyelination: From Aspiration to Reality. Cell Stem Cell, 16, 576–7.

Franklin, R. J. & Goldman, S. A. 2015. Glia Disease and Repair-Remyelination. Cold Spring Harb Perspect Biol, 7, a020594.

Franklin, R. J. M. & Ffrench-Constant, C. 2017. Regenerating CNS myelin - from mechanisms to experimental medicines. Nat Rev Neurosci, 18, 753–769.

Furusho, M., Dupree, J. L., Nave, K. A. & Bansal, R. 2012. Fibroblast growth factor receptor signaling in oligodendrocytes regulates myelin sheath thickness. J Neurosci, 32, 6631–41.

Govier-Cole, A. E., Wood, R. J., Fletcher, J. L., Gonsalvez, D. G., Merlo, D., Cate, H. S., Murray, S. S. & Xiao, J. 2019. Inhibiting Bone Morphogenetic Protein 4 Type I Receptor Signaling Promotes Remyelination by Potentiating Oligodendrocyte Differentiation. eNeuro, 6.

Habuchi, H., Habuchi, O. & Kimata, K. 2004. Sulfation pattern in glycosaminoglycan: does it have a code? Glycoconj J, 21, 47–52.

Harlow, D. E. & Macklin, W. B. 2014. Inhibitors of myelination: ECM changes, CSPGs and PTPs. Exp Neurol, 251, 39–46.

Holst, C. R., Bou-Reslan, H., Gore, B. B., Wong, K., Grant, D., Chalasani, S., Carano, R. A., Frantz, G. D., Tessier-Lavigne, M., Bolon, B., French, D. M. & Ashkenazi, A. 2007. Secreted sulfatases Sulf1 and Sulf2 have overlapping yet essential roles in mouse neonatal survival. PLoS One, 2, e575.

Jenniskens, G. J., Oosterhof, A., Brandwijk, R., Veerkamp, J. H. & Van Kuppevelt, T. H. 2000. Heparan sulfate heterogeneity in skeletal muscle basal lamina: demonstration by phage display-derived antibodies. J Neurosci, 20, 4099–111.

Kakinuma, Y., Saito, F., Ohsawa, S., Furuichi, T. & Miura, M. 2004. A sulfatase regulating the migratory potency of oligodendrocyte progenitor cells through tyrosine phosphorylation of beta-catenin. J Neurosci Res, 77, 653–61.

Karoli, T., Liu, L., Fairweather, J. K., Hammond, E., Li, C. P., Cochran, S., Bergefall, K., Trybala, E., Addison, R. S. & Ferro, V. 2005. Synthesis, biological activity, and preliminary pharmacokinetic evaluation of analogues of a phosphosulfomannan angiogenesis inhibitor (PI-88). J Med Chem, 48, 8229–36.

Keough, M. B., Rogers, J. A., Zhang, P., Jensen, S. K., Stephenson, E. L., Chen, T., Hurlbert, M. G., Lau, L. W., Rawji, K. S., Plemel, J. R., Koch, M., Ling, C. C. & Yong, V. W. 2016. An inhibitor of chondroitin sulfate proteoglycan synthesis promotes central nervous system remyelination. Nat Commun, 7, 11312.

Korchynskyi, O. & Ten Dijke, P. 2002. Identification and functional characterization of distinct critically important bone morphogenetic protein-specific response elements in the Id1 promoter. J Biol Chem, 277, 4883–91.

Kuhlmann, T., Miron, V., Cui, Q., Wegner, C., Antel, J. & Bruck, W. 2008. Differentiation block of oligodendroglial progenitor cells as a cause for remyelination failure in chronic multiple sclerosis. Brain, 131, 1749–58.

Lamanna, W. C., Baldwin, R. J., Padva, M., Kalus, I., Ten Dam, G., Van Kuppevelt, T. H., Gallagher, J. T., Von Figura, K., Dierks, T. & Merry, C. L. 2006. Heparan sulfate 6-O-endosulfatases: discrete in vivo activities and functional co-operativity. Biochem J, 400, 63–73.

Lau, L. W., Cua, R., Keough, M. B., Haylock-Jacobs, S. & Yong, V. W. 2013. Pathophysiology of the brain extracellular matrix: a new target for remyelination. Nat Rev Neurosci, 14, 722–9.

Lortat-Jacob, H., Kleinman, H. K. & Grimaud, J. A. 1991. High-affinity binding of interferon-gamma to a basement membrane complex (matrigel). J Clin Invest, 87, 878–83.

Mei, F., Fancy, S. P., Shen, Y. A., Niu, J., Zhao, C., Presley, B., Miao, E., Lee, S., Mayoral, S. R., Redmond, S. A., Etxeberria, A., Xiao, L., Franklin, R. J., Green, A., Hauser, S. L. & Chan, J. R. 2014. Micropillar arrays as a high-throughput screening platform for therapeutics in multiple sclerosis. Nat Med, 20, 954–60.

Mi, S., Hu, B., Hahm, K., Luo, Y., Kam Hui, E. S., Yuan, Q., Wong, W. M., Wang, L., Su, H., Chu, T. H., Guo, J., Zhang, W., So, K. F., Pepinsky, B., Shao, Z., Graff, C., Garber, E., Jung, V., Wu, E. X. & Wu, W. 2007. LINGO-1 antagonist promotes spinal cord remyelination and axonal integrity in MOG-induced experimental autoimmune encephalomyelitis. Nat Med, 13, 1228–33.

Morimoto-Tomita, M., Uchimura, K., Werb, Z., Hemmerich, S. & Rosen, S. D. 2002. Cloning and characterization of two extracellular heparin-degrading endosulfatases in mice and humans. J Biol Chem, 277, 49175–85.

Najm, F. J., Madhavan, M., Zaremba, A., Shick, E., Karl, R. T., Factor, D. C., Miller, T. E., Nevin, Z. S., Kantor, C., Sargent, A., Quick, K. L., Schlatzer, D. M., Tang, H., Papoian, R., Brimacombe, K. R., Shen, M., Boxer, M. B., Jadhav, A., Robinson, A. P., Podojil, J. R., Miller, S. D., Miller, R. H. & Tesar, P. J. 2015. Drug-based modulation of endogenous stem cells promotes functional remyelination in vivo. Nature, 522, 216–20.

Nawroth, R., Van Zante, A., Cervantes, S., Mcmanus, M., Hebrok, M. & Rosen, S. D. 2007. Extracellular sulfatases, elements of the Wnt signaling pathway, positively regulate growth and tumorigenicity of human pancreatic cancer cells. PLoS One, 2, e392.

Noble, M., Murray, K., Stroobant, P., Waterfield, M. D. & Riddle, P. 1988. Platelet-derived growth factor promotes division and motility and inhibits premature differentiation of the oligodendrocyte/type-2 astrocyte progenitor cell. Nature, 333, 560–2.

Otsuki, S., Hanson, S. R., Miyaki, S., Grogan, S. P., Kinoshita, M., Asahara, H., Wong, C. H. & Lotz, M. K. 2010. Extracellular sulfatases support cartilage homeostasis by regulating BMP and FGF signaling pathways. Proc Natl Acad Sci U S A, 107, 10202–7.

Parish, C. R., Freeman, C., Brown, K. J., Francis, D. J. & Cowden, W. B. 1999. Identification of sulfated oligosaccharide-based inhibitors of tumor growth and metastasis using novel in vitro assays for angiogenesis and heparanase activity. Cancer Res, 59, 3433–41.

Phillips, J. J., Huillard, E., Robinson, A. E., Ward, A., Lum, D. H., Polley, M. Y., Rosen, S. D., Rowitch, D. H. & Werb, Z. 2012. Heparan sulfate sulfatase SULF2 regulates PDGFRalpha signaling and growth in human and mouse malignant glioma. J Clin Invest, 122, 911–22.

Pol, S. U., Lang, J. K., O’Bara, M. A., Cimato, T. R., Mccallion, A. S. & Sim, F. J. 2013. Sox10-MCS5 enhancer dynamically tracks human oligodendrocyte progenitor fate. Exp Neurol, 247, 694–702.

Pol, S. U., Polanco, J. J., Seidman, R. A., O’Bara, M. A., Shayya, H. J., Dietz, K. C. & Sim, F. J. 2017. Network-Based Genomic Analysis of Human Oligodendrocyte Progenitor Differentiation. Stem Cell Reports, 9, 710–723.

Properzi, F., Lin, R., Kwok, J., Naidu, M., Van Kuppevelt, T. H., Ten Dam, G. B., Camargo, L. M., Raha-Chowdhury, R., Furukawa, Y., Mikami, T., Sugahara, K. & Fawcett, J. W. 2008. Heparan sulphate proteoglycans in glia and in the normal and injured CNS: expression of sulphotransferases and changes in sulphation. Eur J Neurosci, 27, 593–604.

Pu, A., Mishra, M. K., Dong, Y., Ghorbanigazar, S., Stephenson, E. L., Rawji, K. S., Silva, C., Kitagawa, H., Sawcer, S. & Yong, V. W. 2020. The glycosyltransferase EXTL2 promotes proteoglycan deposition and injurious neuroinflammation following demyelination. J Neuroinflammation, 17, 220.

Pu, A., Stephenson, E. L. & Yong, V. W. 2018. The extracellular matrix: Focus on oligodendrocyte biology and targeting CSPGs for remyelination therapies. Glia, 66, 1809–1825.

Raff, M. C., Lillien, L. E., Richardson, W. D., Burne, J. F. & Noble, M. D. 1988. Platelet-derived growth factor from astrocytes drives the clock that times oligodendrocyte development in culture. Nature, 333, 562–5.

Rosen, S. D. & Lemjabbar-Alaoui, H. 2010. Sulf-2: an extracellular modulator of cell signaling and a cancer target candidate. Expert Opin Ther Targets, 14, 935–49.

Sabo, J. K., Aumann, T. D., Merlo, D., Kilpatrick, T. J. & Cate, H. S. 2011. Remyelination is altered by bone morphogenic protein signaling in demyelinated lesions. J Neurosci, 31, 4504–10.

Sarrazin, S., Lamanna, W. C. & Esko, J. D. 2011. Heparan sulfate proteoglycans. Cold Spring Harb Perspect Biol, 3.

Seffouh, A., Milz, F., Przybylski, C., Laguri, C., Oosterhof, A., Bourcier, S., Sadir, R., Dutkowski, E., Daniel, R., Van Kuppevelt, T. H., Dierks, T., Lortat-Jacob, H. & Vives, R. R. 2013. HSulf sulfatases catalyze processive and oriented 6-O-desulfation of heparan sulfate that differentially regulates fibroblast growth factor activity. FASEB J, 27, 2431–9.

Sevin, C., Benraiss, A., Van Dam, D., Bonnin, D., Nagels, G., Verot, L., Laurendeau, I., Vidaud, M., Gieselmann, V., Vanier, M., De Deyn, P. P., Aubourg, P. & Cartier, N. 2006. Intracerebral adeno-associated virus-mediated gene transfer in rapidly progressive forms of metachromatic leukodystrophy. Hum Mol Genet, 15, 53–64.

Sherman, L. S. & Back, S. A. 2008. A ‘GAG’ reflex prevents repair of the damaged CNS. Trends Neurosci, 31, 44–52.

Sim, F. J., Lang, J. K., Waldau, B., Roy, N. S., Schwartz, T. E., Pilcher, W. H., Chandross, K. J., Natesan, S., Merrill, J. E. & Goldman, S. A. 2006. Complementary patterns of gene expression by human oligodendrocyte progenitors and their environment predict determinants of progenitor maintenance and differentiation. Ann Neurol, 59, 763–79.

Sim, F. J., Mcclain, C. R., Schanz, S. J., Protack, T. L., Windrem, M. S. & Goldman, S. A. 2011. CD140a identifies a population of highly myelinogenic, migration-competent and efficiently engrafting human oligodendrocyte progenitor cells. Nat Biotechnol, 29, 934–41.

Sim, F. J., Windrem, M. S. & Goldman, S. A. 2009. Fate determination of adult human glial progenitor cells. Neuron Glia Biol, 5, 45–55.

Sim, F. J., Zhao, C., Penderis, J. & Franklin, R. J. 2002. The age-related decrease in CNS remyelination efficiency is attributable to an impairment of both oligodendrocyte progenitor recruitment and differentiation. J Neurosci, 22, 2451–9.

Stringer, S. E., Mayer-Proschel, M., Kalyani, A., Rao, M. & Gallagher, J. T. 1999. Heparin is a unique marker of progenitors in the glial cell lineage. J Biol Chem, 274, 25455–60.

Tang, R. & Rosen, S. D. 2009. Functional consequences of the subdomain organization of the sulfs. J Biol Chem, 284, 21505–14.

Tripathi, A., Volsko, C., Garcia, J. P., Agirre, E., Allan, K. C., Tesar, P. J., Trapp, B. D., Castelo-Branco, G., Sim, F. J. & Dutta, R. 2019. Oligodendrocyte Intrinsic miR-27a Controls Myelination and Remyelination. Cell Rep, 29, 904–919 e9.

Van Horssen, J., Bo, L., Dijkstra, C. D. & De Vries, H. E. 2006. Extensive extracellular matrix depositions in active multiple sclerosis lesions. Neurobiol Dis, 24, 484–91.

Viviano, B. L., Paine-Saunders, S., Gasiunas, N., Gallagher, J. & Saunders, S. 2004. Domain-specific modification of heparan sulfate by Qsulf1 modulates the binding of the bone morphogenetic protein antagonist Noggin. J Biol Chem, 279, 5604–11.

Wang, J., O’Bara, M. A., Pol, S. U. & Sim, F. J. 2013. CD133/CD140a-based isolation of distinct human multipotent neural progenitor cells and oligodendrocyte progenitor cells. Stem Cells Dev, 22, 2121–31.

Wang, S., Ai, X., Freeman, S. D., Pownall, M. E., Lu, Q., Kessler, D. S. & Emerson, C. P., JR. 2004. QSulf1, a heparan sulfate 6-O-endosulfatase, inhibits fibroblast growth factor signaling in mesoderm induction and angiogenesis. Proc Natl Acad Sci U S A, 101, 4833–8.

Welliver, R. R., Polanco, J. J., Seidman, R. A., Sinha, A. K., O’Bara, M. A., Khaku, Z. M., Santiago Gonzalez, D. A., Nishiyama, A., Wess, J., Feltri, M. L., Paez, P. M. & Sim, F. J. 2018. Muscarinic receptor M3R signaling prevents efficient remyelination by human and mouse oligodendrocyte progenitor cells. J Neurosci.

Winkler, S., Stahl, R. C., Carey, D. J. & Bansal, R. 2002. Syndecan-3 and perlecan are differentially expressed by progenitors and mature oligodendrocytes and accumulate in the extracellular matrix. Journal Of Neuroscience Research, 69, 477–487.

Wolswijk, G. 1998. Chronic stage multiple sclerosis lesions contain a relatively quiescent population of oligodendrocyte precursor cells. J Neurosci, 18, 601–9.

Woodruff, R. H., Fruttiger, M., Richardson, W. D. & Franklin, R. J. 2004. Platelet-derived growth factor regulates oligodendrocyte progenitor numbers in adult CNS and their response following CNS demyelination. Mol Cell Neurosci, 25, 252–62.

Yu, G., Gunay, N. S., Linhardt, R. J., Toida, T., Fareed, J., Hoppensteadt, D. A., Shadid, H., Ferro, V., Li, C., Fewings, K., Palermo, M. C. & Podger, D. 2002. Preparation and anticoagulant activity of the phosphosulfomannan PI-88. Eur J Med Chem, 37, 783–91.

Zeisel, A., Munoz-Manchado, A. B., Codeluppi, S., Lonnerberg, P., La Manno, G., Jureus, A., Marques, S., Munguba, H., He, L., Betsholtz, C., Rolny, C., Castelo-Branco, G., Hjerling-Leffler, J. & Linnarsson, S. 2015. Brain structure. Cell types in the mouse cortex and hippocampus revealed by single-cell RNA-seq. Science, 347, 1138–42.

Zhu, X., Hill, R. A., Dietrich, D., Komitova, M., Suzuki, R. & Nishiyama, A. 2011. Age-dependent fate and lineage restriction of single NG2 cells. Development, 138, 745–53.

