## Supplemental Text for "Overcoming the inhibitory microenvironment surrounding oligodendrocyte progenitor cells following demyelination"

**Supplemental Figures**

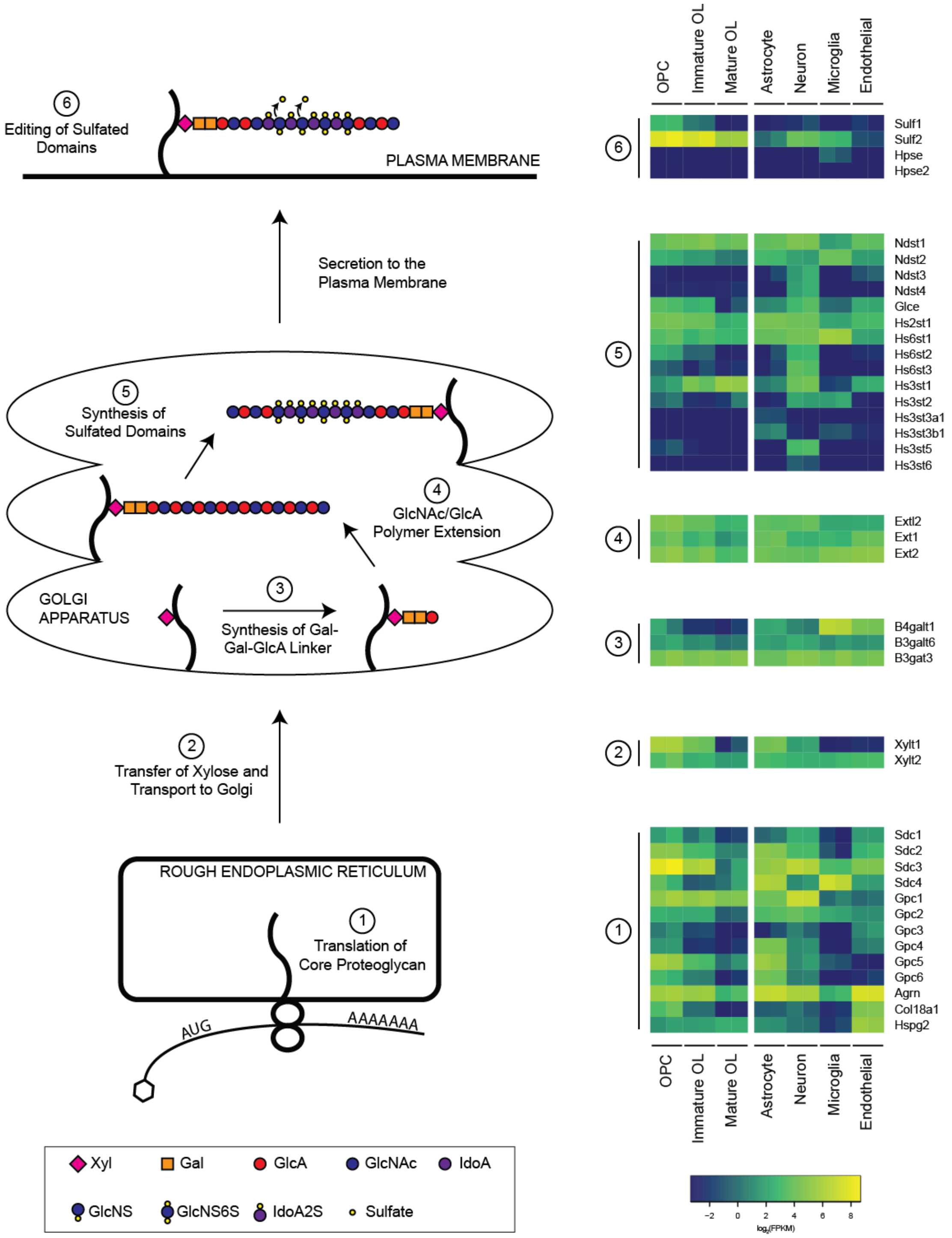

**Figure S1 (related to Figure 1a). Expression of HS-associated genes in the rodent CNS**. The HS biosynthetic pathway is presented as a series of six biological steps beginning with translation of core proteoglycan mRNA in the rough endoplasmic reticulum and concluding with editing of HS side chains at plasma membrane (Esko & Selleck; Fernandez-Vega et al., 2013; Malfait et al., 2013). HS-associated genes have been classified based on the involvement of their protein products in one of these six biosynthetic steps. The heat map presents gene expression on a log(FPKM) scale across murine purified CNS cell populations (Zhang et al., 2014).

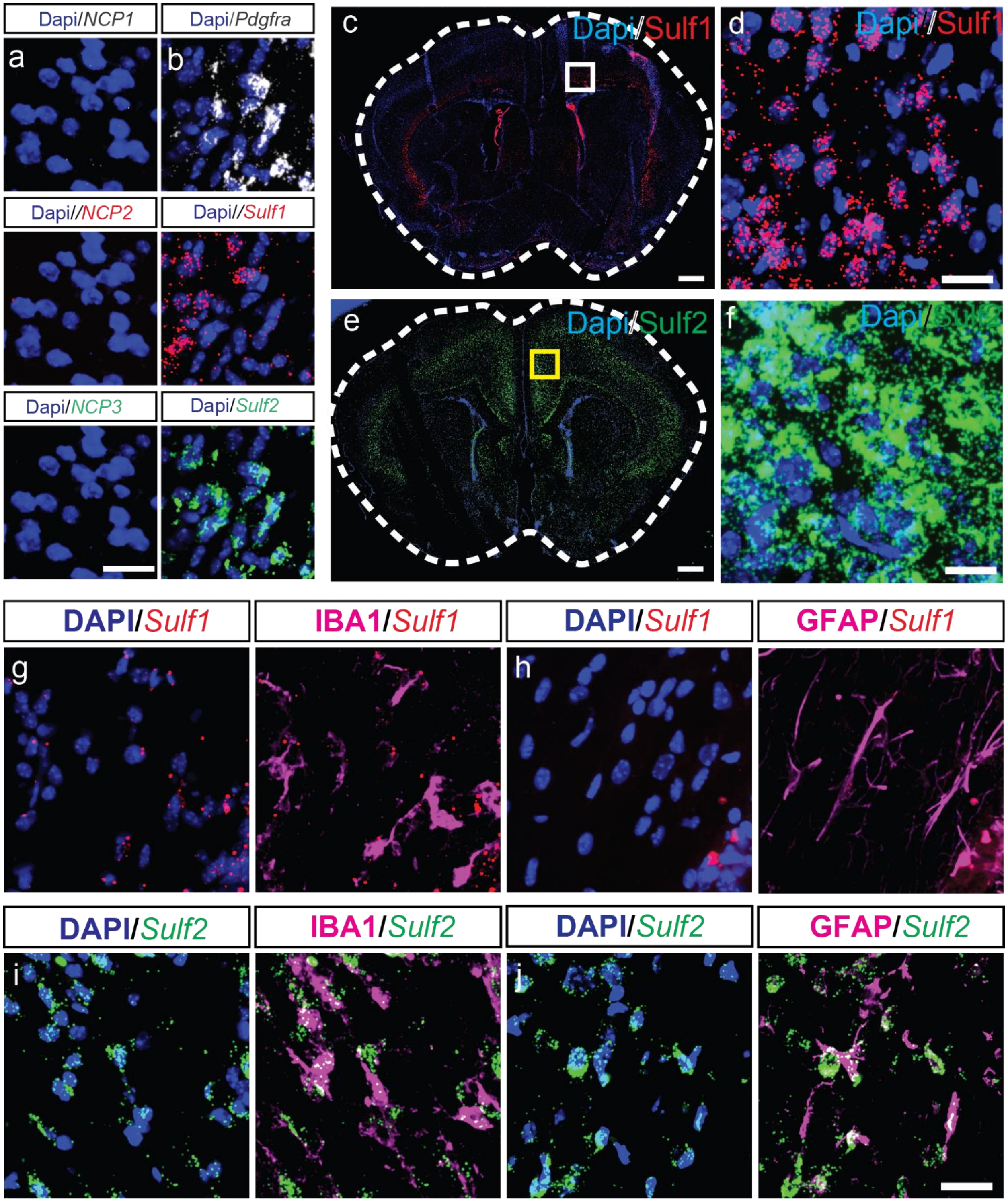

**Figure S2 (related to Figure 1). Sulf1 and 2 expression in postnatal day 7 mouse grey matter and white matter.** **a,** RNAscope *in situ* hybridization (ISH)) of negative control probe 1, 2 and 3 demonstrating the lack of specific binding in mouse brain. **b,** inclusion of *Pdgfra*, *Sulf1* or *Sulf2* probe results in specific and distinct patterns of expression in neonatal white matter. **c-f,** *Sulf1* and *Sulf2* mRNA transcripts are abundant in developing cortical layer V and VI neurons, respectively, and correspond to previously described patterns of expression (Zeisel et al., 2015)(Allen Brain Atlas). **g-j,** RNAscope *in situ* hybridization (ISH) and immunohistochemistry (IHC) of P7 mice brain revealed *Sulf2* expression by a subset of Gfap+ astrocytes and Iba1+ microglia in white matter (both by immunohistochemistry). In contrast, Sulf1 mRNA was not colocalized with Iba1 or and Gfap positive cells. Scale: **a-b,** 20 µm, **c-d**, 100 µm (a) and 20 µm (e-j).

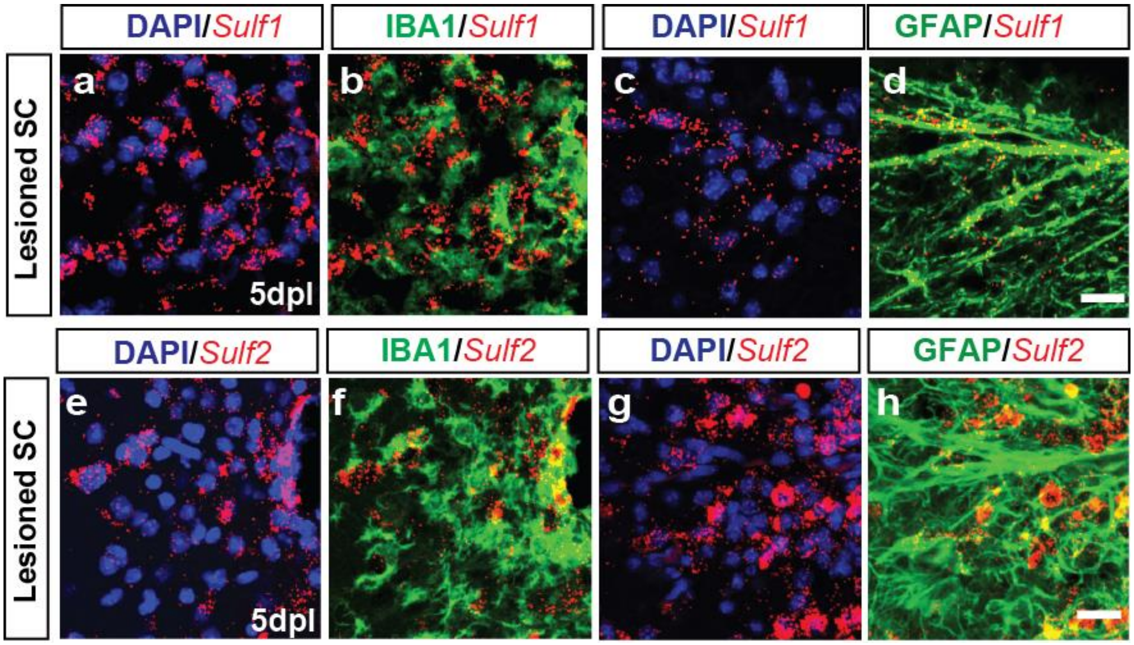

**Figure S3 (related to Figure 2). Increased Sulf2 expression in other glial cells after demyelination in lesion area.** RNAscope *in situ* hybridization (ISH) and immunohistochemistry (IHC) of mice spinal cord after demyelination revealed *Sulf 1* and *2* expression by astrocytes and microglial in lesions as assessed by immunohistochemistry of Gfap and Iba1 and *Sulf1* and *2* mRNA transcripts. Scale: 20 µm.

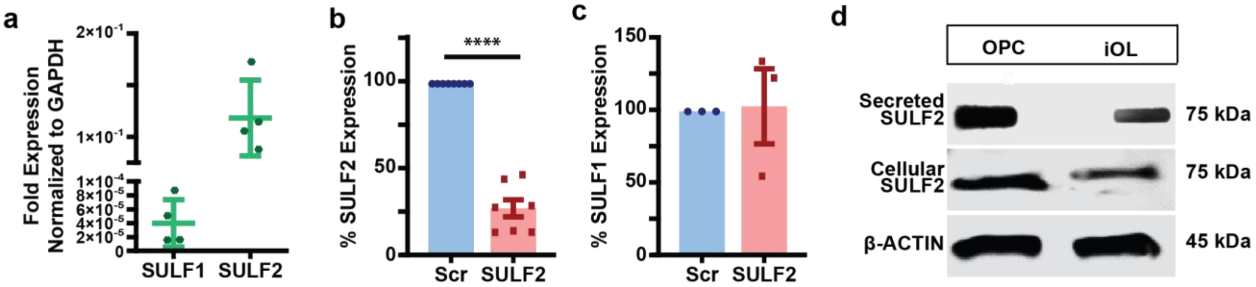

**Figure S4 (related to Figure 3). Lentiviral-mediated expression of SULF2-targeting shRNA reduces *SULF2* mRNA. a,** *Sulf1* and *2*-fold expression in hOPC normalized to GAPDH. **b,** hOPCs transduced with a lentivirus expressing a shRNA targeting SULF2 show significantly reduced expression of SULF2 mRNA, compared to non-targeted scrambled (Scr) shRNA. (****P<0.0001 one sample t test vs 100, n = 8) (**c**) SULF1 expression was not significantly affected by targeted SULF2 knockdown, compared to Scr controls. (P > 0.05 one-sample t test vs 100, n = 3) Graphs represent mean ± SEM normalized to Scr. **d**, Western blot for SULF2 and β-Actin from cultured hOPCs in the presence or absence of growth factors for 3 days to initiate oligodendrocyte differentiation labeled as immature OL (iOL). 30µg protein was examined by slot blot and western blot using anti-Sulf2 antibody. We observed a decrease in secreted and cellular Sulf2 protein levels in iOL as compared to OPC secreted and cellular Sulf2 protein level

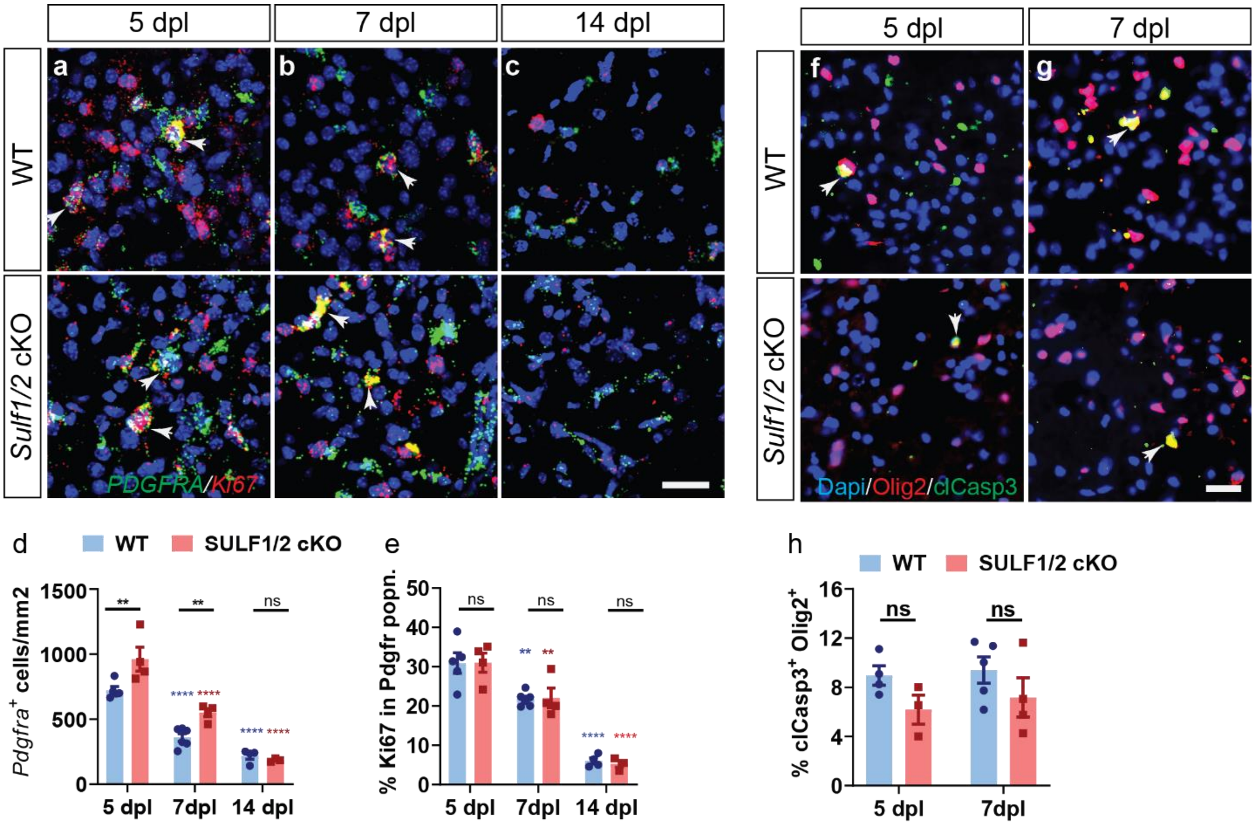

**Figure S5 (related to Figure 4). Conditional ablation of *Sulf1/2* accelerates recruitment of OPCs following demyelination.** Tamoxifen-dependent OPC-specific ablation of both *Sulf1/2* in $NG2^+$ OPCs was initiated prior to demyelination. Control animals (lacking cre) were treated in an identical manner. **a-c,** *Pdgfra*$^+$ cell density and proliferation ($Ki67^+Pdgfra^+$) was assessed at 5, 7, and 14 days post-lesion (dpl) by dual RNAscope *in situ* hybridization (ISH). *Pdgfra* (green), *Ki67* (red) and DAPI (blue). White arrows denote *Pdgfra*$^+$ OPCs that co-expresses *Ki67*. The density (cells/mm$^2$) of $Pdgfra^+$ oligodendrocytes lineage cells (**d**) and percentage of $Pdgfra^+Ki67^+$ proliferating OPCs (**e**) was quantified (n = 3 - 6 mice per group). Two-way ANOVA for different time points. Holm-Sidak post-test vs. wild-type control *, **, ***, *** indicates p < 0.05, 0.01, 0.001, 0.0001 respectively. Blue denotes p-values compared to control wildtype while red denotes p-values compared to *Sulf1/2* cKO group. *Sulf1/2* ablation significantly increased OPC recruitment (**d**) at 5 and 7dpl compared to WT control. While there was no difference in the proportion of proliferating $Ki67^+Pdgfra^+$ cells compared to wild type at each time point. **f-g,** Cell death was assessed by cleaved caspase 3 colocalization with $Olig2^+$ oligodendrocyte lineage cells and percentage of $Olig2^+clCasp3^+$ apoptotic cells of percentage were quantified among total $Olig2^+$ oligodendrocyte lineage cells **(h)** at 5 and 7dpl (n = 3 - 4 mice per group). Mean ± SEM shown. *, **, ***, **** indicate Two-way ANOVA Holm-Sidak post-test p < 0.05, <0.01, <0.001, and < 0.0001 respectively. There was no significant difference in the proportion of apoptotic $clCasp3^+Olig2^+$ cells compared to wild type at each time points Scale: 20 µm.

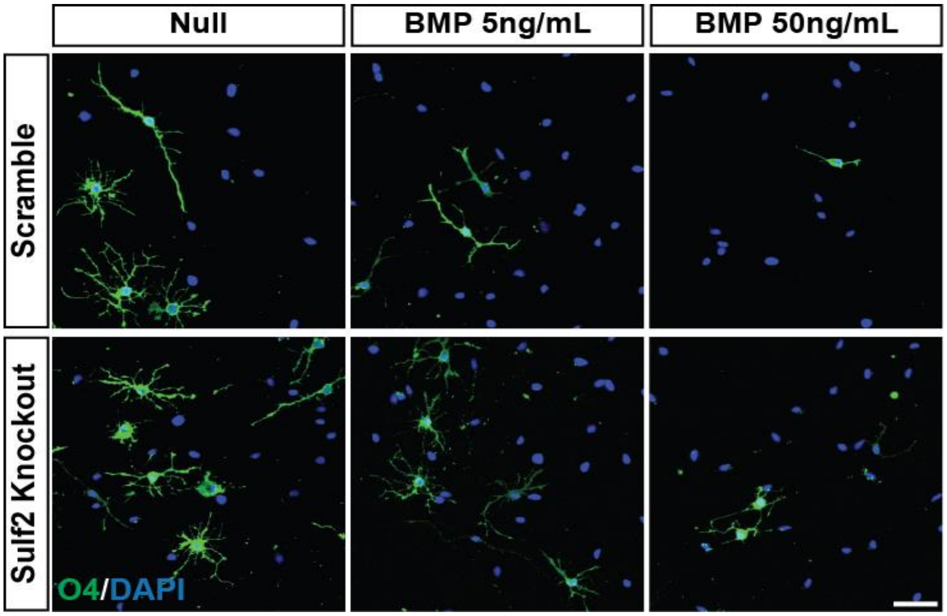

**Figure S6 (related for Figure 5). Sulfatase 2 knockdown increases differentiation despite inhibitory BMP signaling.** Human OPCs were transduced with Sulf2-targeted lentivirus or a scrambled control and allowed to differentiate in the absence or presence of BMP7, as indicated. After four days of differentiation, cultures were live-stained with differentiation marker O4 (green), and DAPI (blue) following fixation. Human OPC differentiation is significantly reduced following BMP7 treatment. SULF2 knockdown attenuated the effects of BMP, significantly increased differentiation of human OPCs, even in the absence of exogenous BMP, compared to scramble knockdown controls. Quantification in Figure 5. Scale: 100µm.

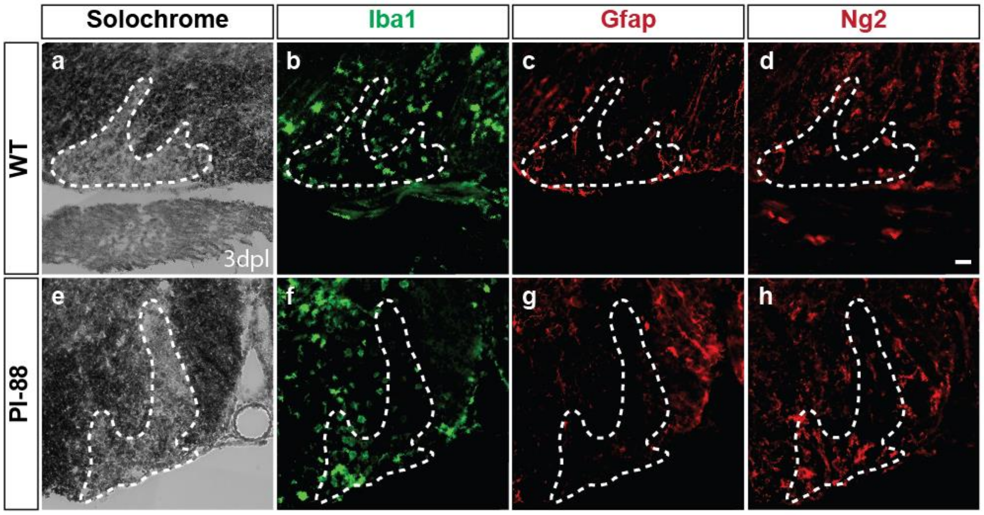

**Figure S7 (related to Figure 7). Administration of PI-88 does not alter lesion dynamics at 3 days post-lesion.** Adult mice were subjected to lysolecithin-mediated focal demyelination of the spinal cord with or without simultaneous administration of 10µg/ml PI-88 directly into the lesion site. Animals were sacrificed at 3 days post-lesion (3dpl) to assess lesion dynamics. Demyelinated lesions of similar size were observed in the ventral white matter of mice (**a-e**) in both experimental groups. (**b-f**) Microglial infiltration, activation and proliferation were similar across experimental groups as assessed by Iba1 immunohistochemistry. (**c-g**) Gliosis was comparable in control and PI-88 treated animals, as assessed by GFAP immunohistochemistry. Oligodendrocyte progenitor cell infiltration was similar in control and experimental groups, as assessed by NG2 immunohistochemistry (**d-h**). n=3--6 mice per group. Scale: 20 µm (**a-h**).

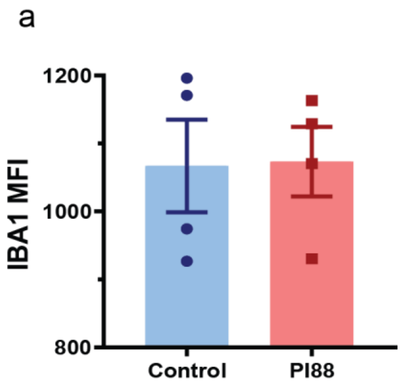

**Figure S8 (related to Figure 7). PI-88 treatment does not increase microglial infiltration in demyelinating lesions.** PI-88 was injected into demyelinated lesions in young adult mouse spinal cord at the time of lysolecithin injection. **a.** Microglial responses was Iba1 immunofluorescence at 7 dpl (n = 3-4 animals, mean ± SEM).

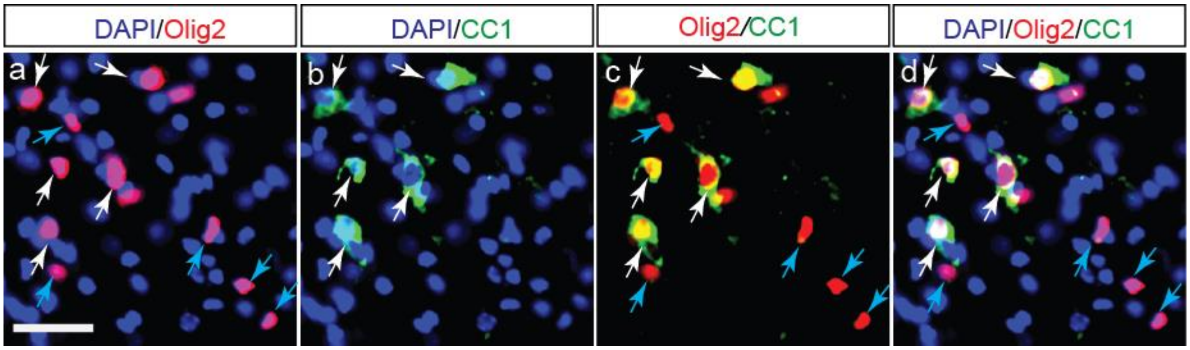

**Figure S9 (related to Methods). Method for quantifying mature oligodendrocytes. a-d.** For identification and quantification of mature oligodendrocytes, we used Olig2/CC1 immunofluorescence and wide-field epifluorescence microscopy. $Olig2^+$ cells (red) with perinuclear $CC1^+$ (green) cells were considered as $Olig2^+CC1^+$ mature oligodendrocytes. White arrows indicate selection of $Olig2^+CC1^+$ oligodendrocytes (b-d). and blue arrow represent $Olig2^+CC1^-$ cells (c-d). Density of mature oligodendrocytes was calculated by counting cells within lesion border and the extent of differentiation assessed by measuring the percentage of $CC1^+$ cells within the Olig2 population. Scale: 60µm.

**Supplemental Tables**

**Table S1 (related to Figure 1) – Comparison of human and mouse HS-related gene expression**

| Human Symbol | Description | Mouse OPC FPKM (FPKM) | Human OPC (FPKM) | Enrichment in OL lineage (Score) |
|---|---|---|---|---|
| SULF1 | Sulfatase 1 | 7.5 ± 0.4 | 0.2 ± 0.1 | 0.92 |
| SULF2 | Sulfatase 2 | 262.5 ± 53.0 | 99.9 ± 8.4 | 0.95 |
| XYLT1 | Xylosyltransferase I | 63 ± 0.5 | 46.2 ± 7.3 | 0.78 |
| HS3ST1 | Heparan sulfate (glucosamine) 3-O-sulfotransferase 1 | 3.1 ± 0.8 | ND | 0.72 (very high in mature OL) |
| EXTL2 | Exostoses (multiple)-like 2 | 26.4 ± 1.9 | 14.6 ± 1.2 | 0.57 |
| CSPG4 | Chondroitin sulfate proteoglycan 4 | 117.4 ± 15.5 | 88.0 ± 2.8 | 0.91 |
| PDGFRA | PDGF receptor, alpha polypeptide | 596.2 ± 32.4 | 422 ± 15.4 | 0.99 |
| SOX10 | SRY-box 10 | 147.1 ± 17.9 | 46.4 ± 0.9 | 0.99 |

For mouse and human FPKM data, mean ± SEM shown, n=2. An oligodendrocyte lineage enrichment score was defined as the sum of expression (FPKM) across the three oligodendrocyte lineage populations (OPC, MOG, new OL) divided by the sum of its expression across all seven CNS populations. Mouse data taken from (Zhang et al., 2014).

**Table S2 (related to Figure 5) – Comparison of effects of WNT and BMP modulating drugs on OPC recruitment and oligodendrogenesis in WT and *Sulf1/2* cKO.**

| | Olig2$^{+}$ cell density (cells / mm$^{2}$) | | CC1$^{+}$ cell density (cells / mm$^{2}$) | | % CC1$^{+}$ cells (among Olig2$^{+}$ cells) | |
|---|---|---|---|---|---|---|
| Condition | WT | *Sulf1/2* cKO | WT | *Sulf1/2* cKO | WT | *Sulf1/2* cKO |
| Control | 655.6 ± 43.7 (n = 8) | 899.7 ± 39.8 (n = 6) | 311.8 ± 37.6 | 718.2 ± 69.4 | 47.8 ± 2.9 | 79.0 ± 5.6 |
| CHIR | 519.3 ± 92.1 (n = 4) | 952.6 ± 45.0 (n = 6) | 111.1 ± 25.8 | 695.8 ± 28.5 | 20.8 ± 2.5 | 73.6 ± 3.4 |
| XAV | 808.9 ± 50.8 (n = 8) | 837.2 ± 63.8 (n = 5) | 624.9 ± 56.4 | 671.5 ± 68.0 | 76.8 ± 3.3 | 80.2 ± 4.9 |
| A01 | 552.7 ± 43.3 (n = 4) | 688.0 ± 54.7 (n = 7) | 106.9 ± 33.8 | 403.6 ± 47.5 | 18.4 ± 4.8 | 54.4 ± 7.4 |
| LDN | 957.7 ± 17.8 (n = 5) | 1147.3 ± 37.1 (n = 3) | 811.2 ± 38.0 | 894.8 ± 71.0 | 85.6 ± 2.4 | 80.6 ± 4.4 |

**Table S3 (related to Figure 5) – Two-way ANOVA showing effects of WNT and BMP modulation in WT and *Sulf1/2* cko and the interaction.**

| | Olig2$^{+}$ cell density | | CC1$^{+}$ cell density | | %CC1$^{+}$ cells | |
|---|---|---|---|---|---|---|
| | F (DFn, DFd) | p value | F (DFn, DFd) | p value | F (DFn, DFd) | p value |
| Interaction | F(4,46)=3.894 | 0.0083 | F(4,46)=8.488 | <0.0001 | F(4,46)=12.23 | <0.0001 |

| | | | | | | |
|---|---|---|---|---|---|---|
| **Drug vs Control** | **F(4,46)=13.91** | **<0.0001** | **F(4,46)=31.08** | **<0.0001** | **F(4,46)=34.17** | **<0.0001** |
| **Wt vs Sulf1/2 cko** | **F(1,46)=35.34** | **<0.0001** | **F(1,46)=67.39** | **<0.0001** | **F(1,46)=64.58** | **<0.0001** |

**Table S4 (related to Figure 5) Holm-sidak's multiple comparison showing effects of WNT and BMP modulation effects on Olig2, CC1 density and percentage differentiation control vs drug in WT and *Sulf1/2* cKO**

| | **Olig2$^+$ cell density** | | **CC1$^+$ cell density** | | **%CC1$^+$ cells** | |
|---|---|---|---|---|---|---|
| **Wild Type** | | | | | | |
| | **Significance** | **Adjusted P value** | **Significance** | **Adjusted P value** | **Significance** | **Adjusted P value** |
| **Control vs CHIR** | ns | 0.2185 | * | 0.0291 | *** | 0.0004 |
| **Control vs XAV-939** | ns | 0.0824 | **** | <0.0001 | **** | <0.0001 |
| **Control vs A01** | ns | 0.3298 | * | 0.0291 | *** | 0.0002 |
| **Control vs LDN-193189** | *** | 0.0008 | **** | <0.0001 | **** | <0.0001 |
| ***Sulf1/2* cKO** | | | | | | |
| **Control vs CHIR** | ns | 0.6506 | ns | 0.9350 | ns | 0.8943 |
| **Control vs XAV-939** | ns | 0.6506 | ns | 0.8981 | ns | 0.9953 |
| **Control vs A01** | * | 0.0246 | *** | 0.0003 | ** | 0.0020 |
| **Control vs LDN-193189** | * | 0.0409 | ns | 0.1785 | ns | 0.9953 |

**Table S5 (related to Figure 5) Holm-sidak's multiple comparison showing effects of WNT and BMP modulation effects on Olig2, CC1 density and percentage differentiation between WT and *Sulf1/2* cko**

| | **Olig2$^+$ cell density** | | **CC1$^+$ cell density** | | **%CC1$^+$ cells** | |
|---|---|---|---|---|---|---|
| **WT vs *Sulf1/2* cko** | **Significance** | **Adjusted P value** | **Significance** | **Adjusted P value** | **Significance** | **Adjusted P value** |
| **Control** | ** | 0.0027 | **** | <0.0001 | **** | <0.0001 |
| **CHIR-99021** | **** | <0.0001 | **** | <0.0001 | **** | <0.0001 |
| **XAV-939** | ns | 0.6904 | ns | 0.5860 | ns | 0.7572 |
| **A01** | ns | 0.1683 | ** | 0.0011 | **** | <0.0001 |
| **LDN-193189** | ns | 0.1198 | ns | 0.5860 | ns | 0.7572 |

**Table S6 (related to Figure 8) – Two-way ANOVA showing effects PI88 treatment in WT and *Sulf1/2* cKO and the interaction.**

| | **Olig2$^+$ cell density** | | **CC1$^+$ cell density** | | **%CC1$^+$ cells** | |
|---|---|---|---|---|---|---|
| | **F (DFn, DFd)** | **p value** | **F (DFn, DFd)** | **p value** | **F (DFn, DFd)** | **p value** |
| **Interaction** | **F(1,13)=8.368** | **<0.05** | **F(1,12)=14.42** | **<0.001** | **F(1,12)=9.993** | **<0.001** |
| **Drug vs Control** | **F(1,13)=0.460** | **0.50** | **F(1,12)=5.147** | **<0.05** | **F(1,12)=11.05** | **<0.05** |
| **Wt vs Sulf1/2 cko** | **F(1,13)=1.93** | **0.18** | **F(1,12)=32.33** | **<0.0001** | **F(1,12)=81.42** | **<0.0001** |

### References

Esko, J. D., & Selleck, S. B. (2002). Order out of chaos: assembly of ligand binding sites in heparan sulfate. *Annu Rev Biochem, 71*(1), 435-471. doi:10.1146/annurev.biochem.71.110601.135458

Fernandez-Vega, I., Garcia, O., Crespo, A., Castanon, S., Menendez, P., Astudillo, A., & Quiros, L. M. (2013). Specific genes involved in synthesis and editing of heparan sulfate proteoglycans show altered expression patterns in breast cancer. *BMC Cancer, 13*, 24. doi:10.1186/1471-2407-13-24

Malfait, F., Kariminejad, A., Van Damme, T., Gauche, C., Syx, D., Merhi-Soussi, F., . . . De Paepe, A. (2013). Defective initiation of glycosaminoglycan synthesis due to B3GALT6 mutations causes a pleiotropic Ehlers-Danlos-syndrome-like connective tissue disorder. *Am J Hum Genet, 92*(6), 935-945. doi:10.1016/j.ajhg.2013.04.016

Zeisel, A., Munoz-Manchado, A. B., Codeluppi, S., Lonnerberg, P., La Manno, G., Jureus, A., . . . Linnarsson, S. (2015). Brain structure. Cell types in the mouse cortex and hippocampus revealed by single-cell RNA-seq. *Science, 347*(6226), 1138-1142. doi:10.1126/science.aaa1934

Zhang, Y., Chen, K., Sloan, S. A., Bennett, M. L., Scholze, A. R., O'Keeffe, S., . . . Wu, J. Q. (2014). An RNA-sequencing transcriptome and splicing database of glia, neurons, and vascular cells of the cerebral cortex. *J Neurosci, 34*(36), 11929-11947. doi:10.1523/JNEUROSCI.1860-14.2014
